# Selective knockout of PKA regulatory subunits reveal opposite catalytic and metabolic consequences with implications for Alzheimer’s disease

**DOI:** 10.64898/2026.06.26.734839

**Authors:** Leigh-Ana M. Rossitto, Tsanwen Lu, Yuliang Ma, Pallavi Kaila Sharma, Valeria Burghi, Carlos C. Gonzalez, Jessica Bruystens, Svetlana Maurya, Jian Wu, Alexis Lona, Irina Kufareva, J. Silvio Gutkind, David J. Gonzalez, Xu Chen, Susan S. Taylor

## Abstract

cAMP-dependent Protein Kinase A (PKA) is a master regulator of cell signaling involved in energy metabolism, synaptic plasticity, and stress response. Dysregulated PKA signaling is implicated in diseases including neurodegeneration and cancer. PKA catalytic activity is regulated by two nonredundant regulatory subunits, Type I (RIα/RIβ) and Type II (RIIα/RIIβ), whose divergent functions are not fully understood. We generated double-knockout (KO) cell lines of RIα/RIβ and RIIα/RIIβ subunits and performed multiplexed MS-based proteomic and phosphoproteomic profiling under basal and glucose-perturbed conditions. We found that RI and RII loss drives distinct, and often opposite, remodeling of the cellular proteome and phosphoproteome. While both mutants blunted metabolic flexibility to glycolytic stressors and stimuli, RI and RII KO cells exhibited elevated and depressed glycolytic signaling, respectively. Interestingly, RI KO increased the abundance and kinase activity of the PKA catalytic subunit Cα isoform, leading to an increase in PKA substrate phosphorylation, whereas RII KO decreased the abundance, kinase activity, and substrate phosphorylation by the catalytic subunit Cβ isoform. Notably, one of the most differentially affected PKA sites between RI and RII KOs maps to Tau, whose hyperphosphorylation is a hallmark of Alzheimer’s disease. Loss of RI increased Tau phosphorylation, which was not only caused by increased PKA catalytic activity, but also a higher binding affinity of Tau to RII subunits on the negatively-charged flexible linker region. Overall, the present study demonstrates that PKA RI and RII subunits play nonredundant roles in modulating PKA activity, metabolic flexibility, and phospho-regulation of key disease-associated substrates such as Tau.

## INTRODUCTION

Protein Kinase A (PKA), or cAMP-dependent protein kinase, is a master regulator of cell signaling. It is found ubiquitously in mammalian cells and is important in numerous biological functions including glucose, glycogen, and lipid metabolism, synaptic plasticity, autophagy, and response to cellular stress [1]. Canonical PKA signaling begins with the activation of transmembrane G Protein-Coupled-Receptors (GPCRs) by extracellular ligands, causing the release of the activated (GTP-bound) form of the cAMP-stimulatory Gα_S_ subunit. The GTP-bound Gα_S_ activates adenylyl cyclase (AC), which converts ATP into cAMP. cAMP can then bind PKA regulatory (R) subunits, releasing them from the catalytic (C) subunits, which can then catalyze the reaction of ATP and a substrate protein to ADP and its substrate phosphoprotein [2]. PKA preferentially phosphorylates substrates containing basic residues (arginine or lysine) upstream of the phosphorylation site (e.g., RRxS). Over 200 substrates have been identified including key regulators of glycolysis and gluconeogenesis such as phosphofructokinase (PFK) and microtubule-associated protein Tau (MAPT), implicated in Alzheimer’s disease and related dementias (ADRD) [3–6].

PKA is a tetrameric holoenzyme with two C subunits and two R subunits (R_2_:C_2_) in its inactive conformation. The C subunits primarily consist of their isoforms Cα and Cβ, encoded by genes *PRKACA* and *PRKACB*, respectively [2], whose functional differences have been extensively explored [7, 8]. In addition, PKA has four R subunit isoforms: RIα and RIβ (Type I) and RIIα and RIIβ (Type II), encoded by *PRKAR1A*, *PRKAR1B*, *PRKAR2A*, and *PRKAR2B*, respectively. The R subunits are similar in sequence and domain organization, with four primary regions: an inhibitor sequence (IS) in the flexible linker which binds to the C subunit active site, two cAMP binding domains (CNB-A and B), and a dimerization/docking (D/D) domain which is a dual site for R_2_ dimer formation and A Kinase Anchoring Proteins (AKAPs) docking. When cAMP binds to the R subunits, they dissociate from the C subunits, which are then free to act on substrates, often from signaling scaffolds created by AKAPs in various subcellular locations [2, 9].

Despite similarities, the four R subunits are considered functionally nonredundant. Biochemically, they contribute to different holoenzyme quaternary structures, they have varying sensitivities to cAMP, and, while RIIα and RIIβ have serine substrate sites in their inhibitor sequences, RIα and RIβ have alanine and glycine pseudosubstrate sites, respectively [10–14]. They are also differentially expressed across tissues: while each is detected in all tissues, RIα has low tissue specificity, RIIα is enriched in skeletal muscle and testis, RIβ is enriched in brain, and RIIβ is enriched in adipose tissue and brain [15]. Additionally, various AKAPs have R subunit specificity, resulting in distinct subcellular localization of different holoenzymes [9]. Studies in murine models have begun to address gaps in knowledge of the distinct functional outcomes [16]. Somatic single-knockouts have drastically different consequences, with RIα loss causing embryonic lethality, RIβ loss causing memory deficits, and RIIβ loss causing motor deficits [17–21]. Mutations in different R subunits are linked to various diseases in humans including cancer, autism, and neurodegenerative disease [1, 22–25]. The molecular underpinning of the divergent functions of the R subunits remains unclear [26, 27].

Despite being one of the most researched kinases in biology, to our knowledge, no study has performed side-by-side comparisons of loss-of-function mutations across all R subunits. The goal of the present study is to elucidate functional nonredundancy between PKA Type I and Type II R subunits. To this end, we employed multiplexed tandem MS-based proteomics and phosphoproteomics, complemented by orthogonal functional and biochemical assays. Overall, we found that loss of RI and RII subunits caused distinct, and often opposite, metabolic and catalytic consequences, including differential effects on C subunit abundance. Moreover, we found that RI and RII KOs differentially regulate phosphorylation of disease-associated protein substrates such as Tau, thereby implicating PKA-R subunit dysfunction in ADRD and unveiling a new connection between metabolic dysregulation and Tau hyperphosphorylation.

## RESULTS

### Loss of PKA Type I versus Type II R subunits shifts the cellular proteome and phosphoproteome in distinct, and often opposite, directions

We employed CRISPR-Cas9 technology to generate Type I (RIα and RIβ) and Type II (RIIα and RIIβ) double-knockout (KO) HEK293T cell lines, hence RI KO and RII KO, respectively, with a scramble guide RNA line as a wild-type (WT) PKA control (**Fig. 1A-B**). HEK cells are one of the few cell lines or types that express all four R subunits endogenously, allowing for side-by-side comparison of individual KOs in the same cellular context (**Fig. S1**) [28]. Sanger sequencing and Western blot (WB) validated the KOs in the single clones (**Fig. 1C**). Next, we performed multiplexed tandem MS to unbiasedly interrogate how each KO alters the global proteomic and phosphoproteomic landscapes under basal and glucose-perturbed conditions. The successful KO was further confirmed by MS (**Fig. 1D-E**), orthogonally validating on-target editing. 8,885 proteins were identified and quantified in our proteomic study, and 2,436 phosphopeptides from 1,223 unique proteins were identified through phosphoproteomics (**Fig. S2**). The abundances of the phosphopeptides were normalized to the total protein abundance unless otherwise specified. We primarily detected phospho-serine (pSer) residues (90.9%), and most detected phosphopeptides (83.4%) contained a single phosphorylated residue (**Fig. S2**). We detected 23 published PKA-C phosphosites, and an additional 25% of the detected sites are predicted PKA-C sites based on motif analysis (**Fig. S2**) [3, 29].

**Figure 1.**
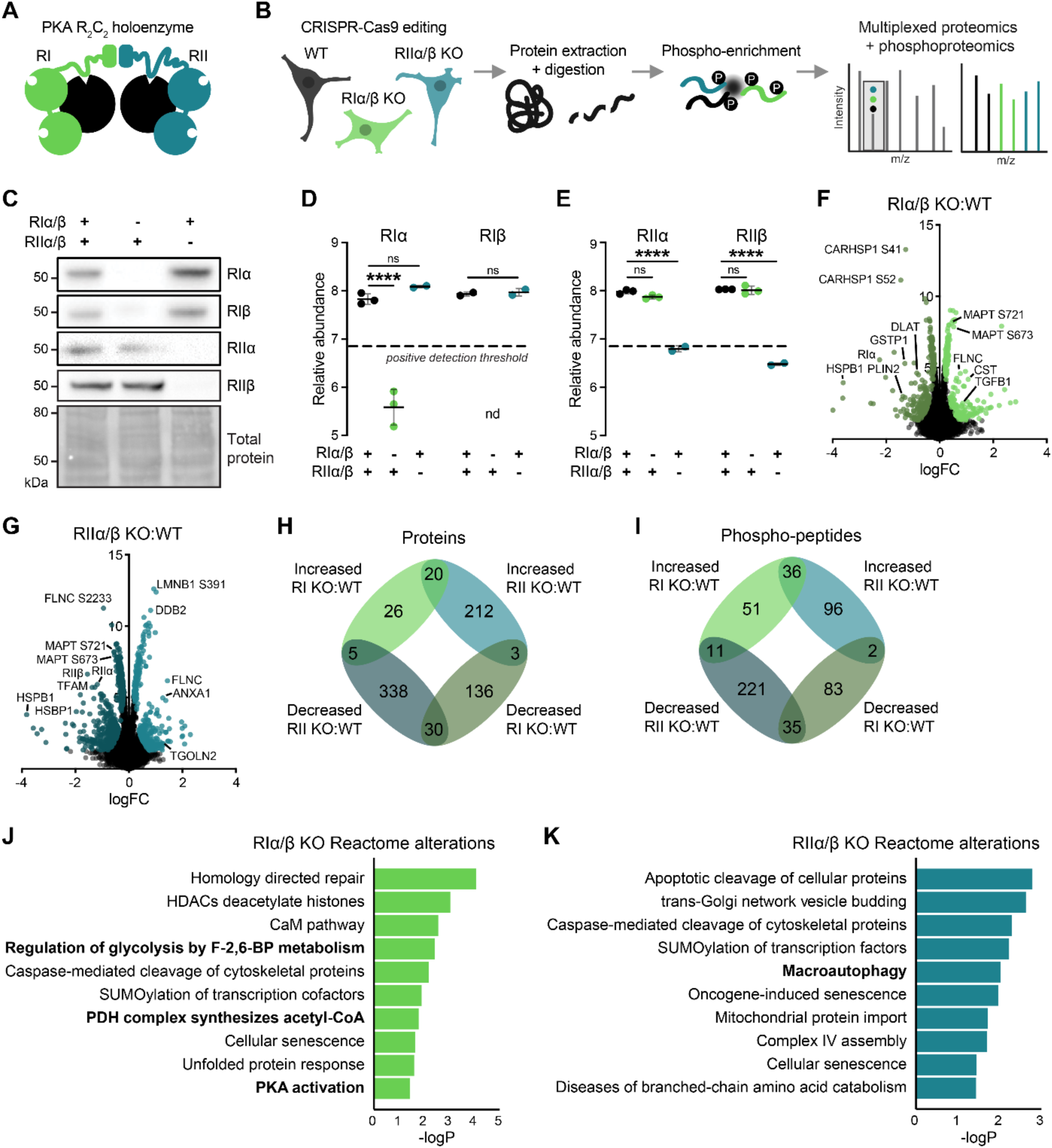
PKA RI and RII double-KO cells show distinct proteomic and phosphoproteomic changes compared to WT HEK293T cells. **(A)** Schematic of RI vs. RII PKA holoenzymes. **(B)** Schematic of MS-based proteomic and phosphoproteomic strategy on PKA RI vs. RII double-KO HEK293T cells. **(C)** Representative WB confirmation of R subunit KOs. **(D-E)** MS quantification of RI (D) and RII subunits (E). Each point represents one independent replicate. The dotted line represents the positive detection threshold, set to 1.5^th^ percentile of all detected protein abundances. nd: not detected. **(F-G)** Volcano plots illustrating binary comparisons between RI KO vs. WT (F) and RII KO vs. WT (G), with the colored dots indicating significantly differentially abundant proteins and phosphosites. **(H-I)** Venn diagrams illustrating shared and unique hits between differentially abundant proteins (H) and phosphosites (I) in RI KO vs. WT and RII KO vs. WT binary comparisons. **(J-K)** Reactome pathway enrichment analysis of differentially abundant total and phosphoproteins for RI KO vs. WT (J) and RII KO vs. WT (K). Data are represented as mean ± SD. P-values for D-G were calculated by limma. Significant hits included in F-K defined as p < 0.05 and π-score, or -logP x |FC| ≥ 1. FC: fold-change.

First, we performed binary comparisons between RI KO versus (vs.) WT and RII KO vs. WT under basal conditions. Each KO had dramatically altered proteomes and phosphoproteomes, but there were surprisingly few overlapping changes between RI KO vs. WT and RII KO vs. WT (**Fig. 1F-I**). Reactome overrepresentation analysis on these hits showed that both mutants affected proteins in cellular stress pathways such as “Caspase-mediated cleavage of cytoskeletal proteins,” “Cellular senescence,” and the broad pathway “Metabolism” (77 & 162 proteins, respectively, representing ∼23% of all hits for either KO) (**Fig. 1J-K**). However, only RI KO but not RII KO affected “PKA activation” (**Fig. 1J-K**). While both KO lines were associated with “energy metabolic dysfunction,” RI KO appeared to specifically affect early glucose metabolism, whereas RII KO affected the electron transport chain and overall mitochondrial function (**Fig. 1J-K**).

### Loss of PKA Type I versus Type II R subunits reveals distinct responses to glycolytic stressors and stimuli

PKA is a key regulator of the balance between glycolysis and gluconeogenesis, responding to glucose availability by phosphorylating central enzymes including PFK-2, thereby directing glycolysis, gluconeogenesis, and glycogen synthesis [1, 5]. To further explore R subunit-specific regulation of glucose metabolism, we mapped all significant changes onto glycolysis/gluconeogenesis (**Fig. S3A**) and the citric acid (TCA) cycle (**Fig. S3B**) at the proteomic and phosphoproteomic levels and identified 41 significant changes in total by p-value (<0.05). Notably, we found differential abundances of the same PKA phosphosite in two PFK-2/FBPase-2 isozymes: PFKFBP2 pS466 decrease with RII KO, and PFKFBP3 pS461 increase with RI KO and decrease with RII KO (**Fig. S3B**). PFK-2/FBPase-2 is a bi-lobal protein that can act as a switch between glycolysis and gluconeogenesis via the interconversion of fructose-6-phosphate and fructose-2,6-bis-phosphate. Activation of its biphosphatase domain (which contains residues S466 and S461 in isozymes B2 and B3, respectively) via phosphorylation results in a shift to gluconeogenesis via PFK-1 inhibition. These findings suggest that RI KO cells exhibit a phosphoproteomic signature consistent with increased gluconeogenic signaling, whereas RII KO cells display a signature consistent with increased glycolytic signaling.

We hypothesized that PKA-R KOs would exhibit disrupted responses to glucose starvation and would not recover with glucose refeeding. To evaluate this, we performed MS-based phosphoproteomics on the cell lines (1) under basal conditions, (2) after glucose starvation (“Stv”) for either 10’ or 30’, and (3) after glucose refeeding for 30’ following 30’ Stv (**Fig. 2A**). Phosphosite profiles were *k*-means clustered to understand how the acute response to glycolytic stress differs in PKA-R KOs compared to WT responses. When applied to the WT data, to identify expected signaling responses to starvation and refeeding, we identified 8 distinct clusters, or patterns of response, via the elbow method (**Fig. 2B**). As shown by the average scaled abundances of the proteins and phosphopeptides within the same clusters, the pattern of responses to starvation and refeeding within the WT cells were blunted in the KO cells (**Fig. 2C**). These data are consistent with the critical role of PKA in regulating glucose metabolism: during starvation, PKA activity decreases significantly, shifting the cell away from glycolysis and energy consumption towards energy preservation, survival, and autophagy. The only two clusters that show increased phosphorylation after 10’ of starvation in WT cells, clusters 5 and 8, are preserved in KO cells, potentially representing PKA-independent stress responses (**Fig. 2C**).

**Figure 2.**
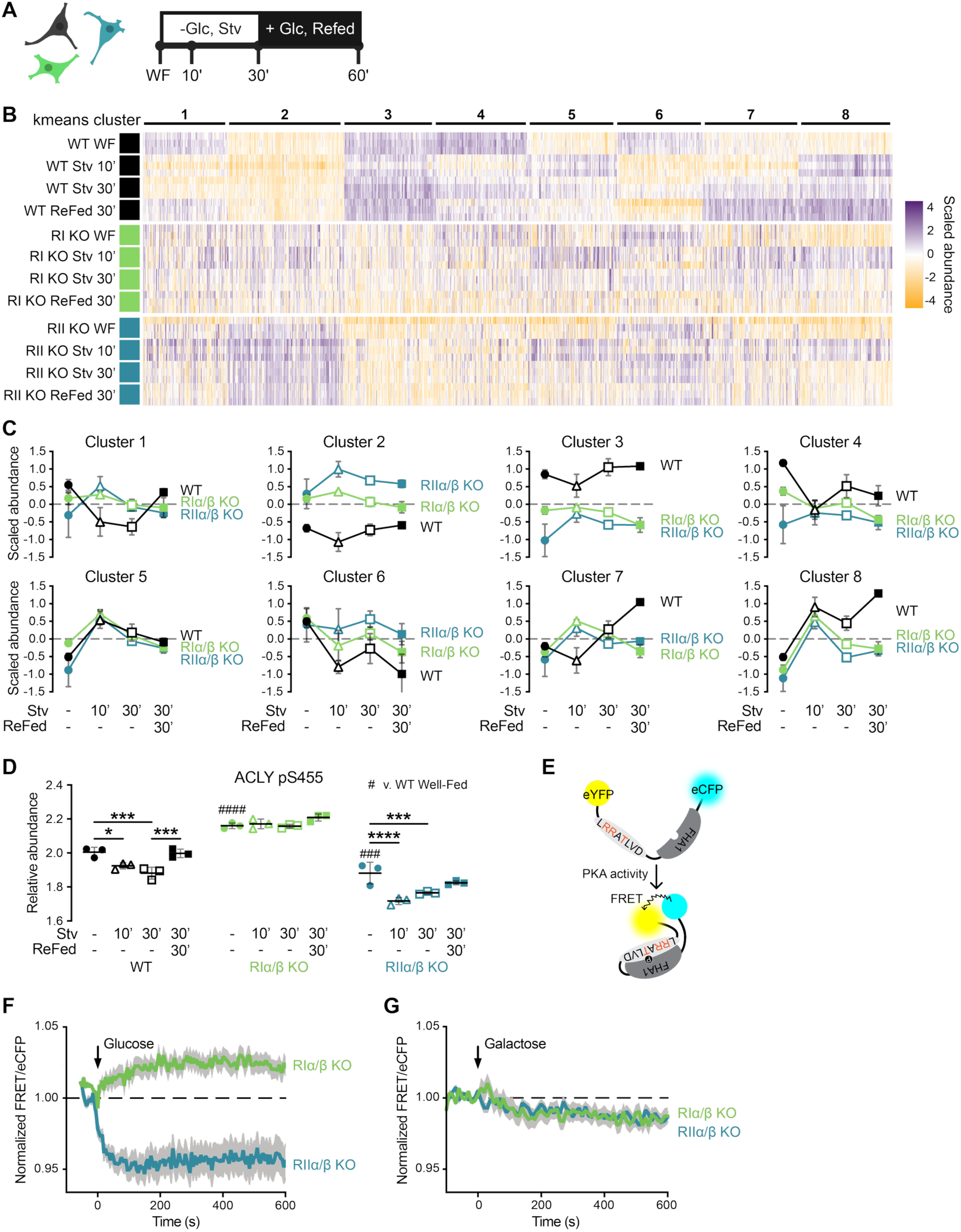
Knockout of PKA-R subunits reveal distinct responses to glucose perturbations. **(A)** Schematic of glucose (glc) starvation (stv) and refeeding paradigm. WF: well-fed (basal). **(B-C)** *k-*means clustering analysis. Heatmap of scaled abundances of phosphopeptides (B) during starvation paradigm, clustered by *k-*means algorithm trained on WT data. Corresponding line graphs (C) depicting group means ± SD. n=3 independent replicates per group. **(D)** Relative abundance ACLY pS455 across starvation and refeeding paradigm. Phosphopeptide abundances were normalized to group average protein abundance. P-values calculated by limma. Comparisons performed: WT vs. RI WF; WT vs. RII WF; within each line, WF vs. Stv 10’, WF vs. Stv 30’, and Stv 30’ vs. Refed 30’. Data are represented as mean ± SD. **(E-G)** PKA activation FRET assay. FRET biosensor design (E). FRET readings upon glucose (F) and galactose (G) bolus, normalized to treatment time at 0 s. Cells were glucose-starved for 4 h prior to treatments. Lines represent mean of n=8-9 replicates per group, and gray shading represents ± SEM.

To delve further into how PKA finely tunes energy metabolism during glucose starvation, we focused on ATP-citrate lyase (ACLY), a key enzyme that diverts citrate away from the TCA cycle to produce cytosolic acetyl-CoA and drive fatty acid synthesis. Phosphorylation of ACLY by PKA at S455 stimulates its enzymatic activity [3, 30]. In WT cells, ACLY pS455 decreased during starvation, indicating a shift towards TCA (**Fig. 2D**). However, ACLY pS455 was elevated in RI KO cells at baseline, consistent with increased PKA activity, and was entirely insensitive to glucose perturbations. In contrast, RII KO cells downregulated pS455 upon glucose withdrawal, while showing a change in recovery dynamics after glucose refeeding compared to WT cells (**Fig. 2D**).

To understand how PKA-R KOs respond to glycolytic stimuli, i.e., glucose and galactose, we treated starved RI and RII KO cells with boluses of each metabolite and measured PKA activation via a reporter that produces a FRET signal upon phosphorylation of the PKA motif in its linker region (**Fig. 2E**). Because galactose does not support glycolytic ATP production to the same extent as glucose, it can be used to distinguish glycolysis-dependent from mitochondrial effects. We found that glucose, but not galactose, treatment differentially altered PKA activity in RI and RII KO cells: RI KO increased PKA activity relative to baseline, whereas RII KO decreased relative activity (**Fig. 2F-G**). In summary, loss of PKA RI or RII resulted in distinct patterns of dysregulated response to glycolytic stimuli and stressors, both leading to failure of the cellular adaptation to glucose perturbations.

### Loss of PKA Type I versus Type II R subunits reveals opposite alterations to catalytic subunit abundance and consequently opposite catalytic output

Reactome analysis of RI KO hits also showed altered “PKA activation,” so we next asked if KO of RI or RII altered basal abundance of PKA-C subunits. Previous studies have reported altered PKA-C activity with RIα loss, although the abundance of the C subunit has not been assessed [17]. MS showed that RI KO significantly increased PKA Cα abundance compared to WT, whereas RII KO significantly decreased Cβ abundance (**Fig. 3A**), both orthogonally validated by WB (**Fig. 3B**). We then asked if the alteration of PKA-C subunit abundance impacts PKA activity using various functional readouts and measures of PKA-C target phosphorylation (**Fig. 3C-H**). PKA-C preferentially binds to and phosphorylates Ser/Thr residues two or three positions following basic residues (RRxS/T). WB probing for pan-PKA substrate phosphorylation (RRxSp/Tp) confirmed that RI KO cells have increased PKA substrate phosphorylation (**Fig. 3C**).

**Figure 3.**
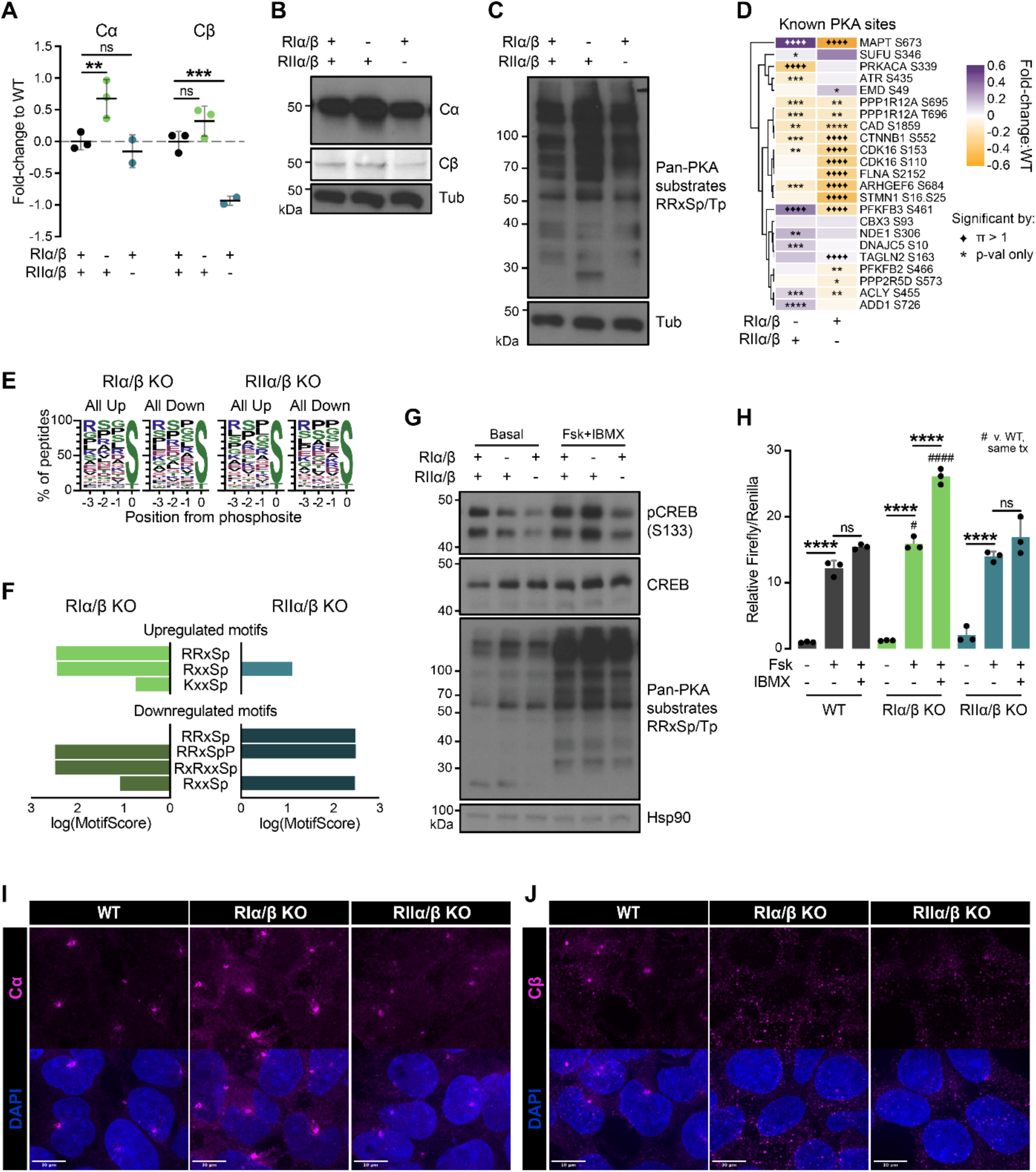
Knockout of PKA-R subunits reveals opposite catalytic consequences. **(A)** MS quantification of PKA-C subunit isoforms by KO cell line, relative to WT. P-values calculated by limma. Data are represented as mean ± SD. **(B)** Representative WB confirmation of PKA-C isoform abundance changes. **(C)** Representative WB of pan-PKA substrates (RRxSp/RRxTp). **(D)** Phosphosite abundances of all known detected PKA sites (n=23) by KO line, relative to WT. P-values calculated by limma. Changes that were additionally significant by π-score, or -logP x |FC| ≥ 1, are denoted with diamonds. FC: fold-change. **(E)** WebLogo depiction of phosphosites from all differentially abundant phosphopeptides (by p-value) in KO vs. WT binary comparisons. Up indicates peptides with significantly increased abundance, and Down indicates peptides with significantly decreased abundance. **(F)** Motif enrichment analysis of differentially abundant peptides (by p-value) in KO vs. WT binary comparisons. Only PKA motifs are shown. Cut-off for significantly enriched motifs was set to p < 0.00001 and minimum number of sequences per motif to 20. **(G)** Representative WB of PKA substrate CREB pS133 and pan-PKA substrates basally and with forskolin (Fsk) and IMBX co-treatment. Both drugs increase intracellular cAMP levels. **(H)** CRE-controlled Firefly luciferase assay to measure basal and maximal PKA activation of each line during Fsk/IBMX treatments. n=3 independent replicates/group. Data are represented as mean ± SD. P-values calculated by Sidák’s multiple comparisons test. All performed comparisons are shown. **(I-J)** Representative confocal imaging of immunofluorescent staining for basal levels of Cα (I) and Cβ (J) in each cell line. Scale bars: 10 μm.

Next, we analyzed known/published PKA sites detected in our phosphoproteomics study (n=23 of 366 sites with known kinases, n=10 sites only documented to be phosphorylated by PKA) [3]. Most of these sites were differentially abundant in binary comparisons between RI or RII KO and WT cells (22/23 sites), with 7 sites increasing and 8 decreasing for RI KO and 2 increasing and 14 decreasing for RII KO (**Fig. 3D**). Beside previously discussed changes in PFK-2/FBPase and ACYL phosphorylation (**Fig. 3D**), we also found that RI KO decreases PKA Cα autophosphorylation at S339 (**Fig. 3D**). Notably, the unnormalized Cα pS339 peptide is still decreased in RI compared to WT (**Table S1**), so this alteration is not entirely due to increased total Cα protein (**Fig. 3A**). S339 (or S338 if excluding the first Met) is in the C-terminal tail of Cα, and while its phosphorylation is not necessary for PKA activation or holoenzyme formation, it is critical for kinase maturation [31]. This finding further illustrates the dysregulation of PKA-C with loss of R subunits, especially RI, although the specific functional outcome of increased total Cα paired with decreased Cα pS339 on Cα is unclear.

Next, we probed our phosphoproteomics data in an unbiased manner via motif enrichment analysis of differentially abundant phosphopeptides to see if PKA motifs were overrepresented in either direction (**Fig. 3E-F**) [32]. Higher motif scores correspond to more specific and statistically significant motifs. PKA motifs were enriched in the list of phosphopeptides with significantly decreased abundance in RII KO cells compared to WT, in line with decreased PKA-driven phosphorylation in RII KO cells, whereas PKA motifs were enriched in both lists of significantly increased and decreased phosphopeptides in RI KO:WT cells (**Fig. 3F**).

When activated, PKA-C translocates to the nucleus to phosphorylate cAMP response element (CRE)-binding protein, CREB [33]. Therefore, we next probed for CREB pS133 under basal and maximal PKA activation conditions by co-treating with AC activator forskolin (Fsk) and phosphodiesterase inhibitor IBMX, both of which increase cAMP levels (**Fig. 3G**). Co-treatment with Fsk and IBMX led to higher levels of pCREB in the RI KO cells as compared to WT and lower in RII KO cells, and changes in pan-PKA substrate (RRxSp/Tp) phosphorylation followed the same pattern (**Fig. 3G**). Next, we measured CRE activation directly via a CRE-controlled Firefly luciferase assay. Consistent with pCREB results, while there were no differences in basal relative Firefly luciferase luminescence, treatment with Fsk or Fsk and IMBX significantly increased such luminescence, and hence CREB activity, in the RI KO, but not the RII KO, compared to WT (**Fig. 3H**).

Lastly, we assessed how the loss of RI or RII affects the subcellular localization of Cα and Cβ via immunofluorescent staining and confocal imaging (**Fig. 3I-J**). Loss of RI or RII alone did not appear to affect Cα localization, although Cα protein abundance was visually changed, consistent with our MS and activity results. However, loss of either RI or RII altered Cβ localization to become more diffuse with smaller puncta (**Fig. 3I-J**). These data together support that loss of RI increases, whereas loss of RII decreases, PKA-C subunit abundance and activity, albeit with substrate-specific exceptions.

### Loss of PKA Type I versus Type II R subunits differentially impacts Tau phosphorylation

Notably, the most differentially abundant known PKA-C phosphosite detected in our study was MAPT pS673, corresponding to pS356 in mature 2N4R Tau (full-length Tau, also known as Tau441). Tau pS356, which is present in all six isoforms of Tau at the fourth repeat domain (RD), was significantly increased with RI loss and decreased with RII loss (**Fig. 1F-G, 3D, 4A**). PKA is one of many known kinases to target Tau, whose hyperphosphorylation is a key pathological event in ADRD, leading to toxic Tau accumulation and formation of neurofibrillary tangles (NFTs). An additional Tau phosphorylation site S404, a well-known early biomarker of ADRD, showed a matching phosphorylation pattern (**Fig. 4B**), consistent with previous reports that PKA-mediated phosphorylation primes the primary Tau kinase GSK-3β to phosphorylate several Tau phosphosites, including S404 [34, 35]. Neither of these changes were driven by changes in total Tau protein abundance (**Fig. S4**). Next, we tracked pS356 with glucose starvation and refeeding and observed that short-term glucose starvation reduced Tau phosphorylation in WT cells (**Fig. 4C**). Glucose starvation has little effect on Tau phosphorylation in RII KO cells, in which pS356 is low basally and remains low with glucose manipulation (**Fig. 4C**). Intriguingly, RI KO cells show a similar dynamic response to the WT cells of pS356 with glucose manipulation, albeit from a significantly higher baseline of phosphorylation (**Fig. 4C**).

**Figure 4.**
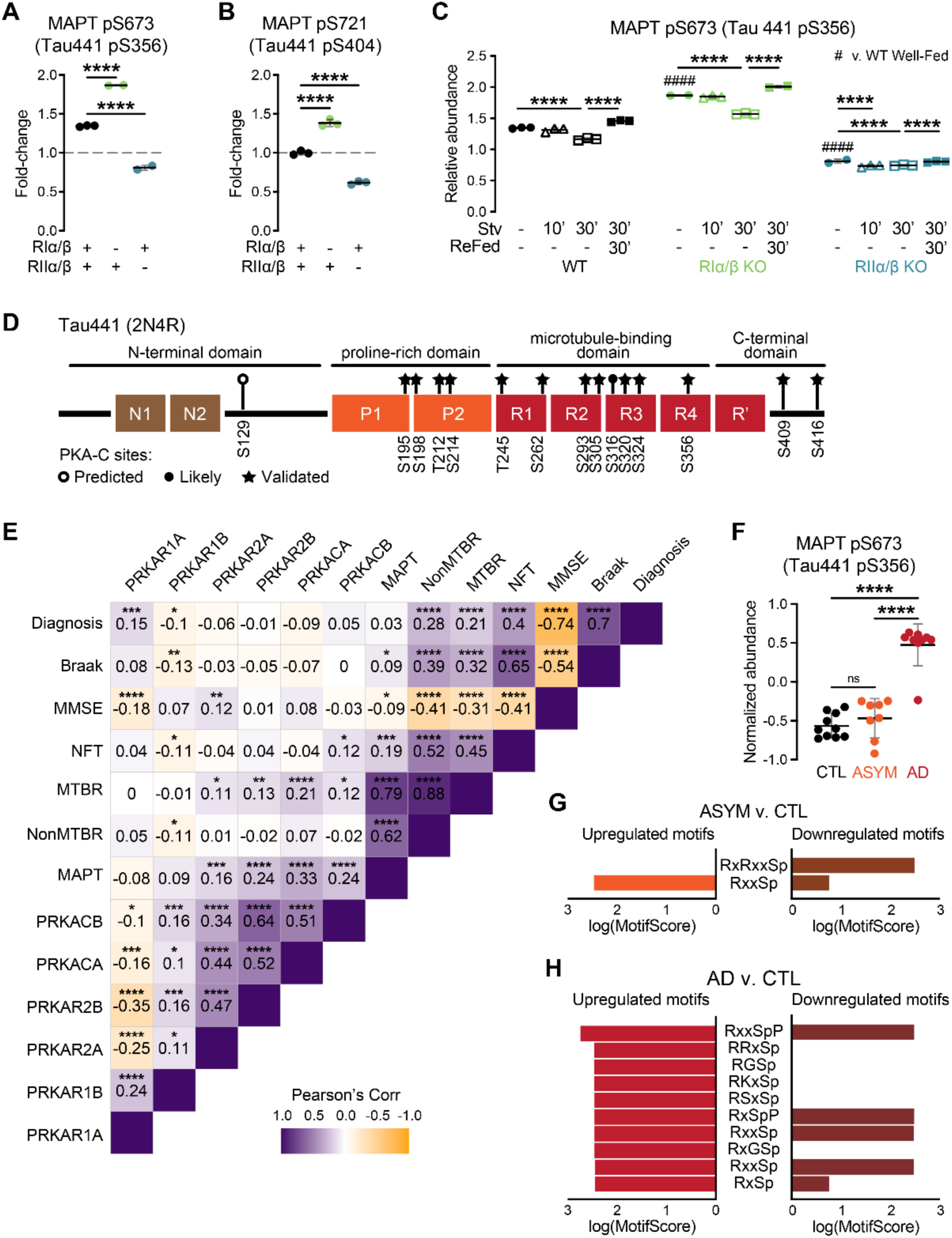
KO of PKA-R subunits differentially impacts Tau phosphorylation. **(A-B)** MS quantification of MAPT PKA phosphosite pS673, corresponding to pS356 in Tau441 (A), and MAPT GSK3β phosphosite pS721, corresponding to pS404 in Tau441 (B), by KO cell line, relative to WT. Phosphopeptide abundances were normalized to group average MAPT abundance. P-values calculated by limma. Data are represented as mean ± SD. **(C)** Relative abundance MAPT PKA phosphosite pS673, corresponding to pS356 in Tau441, across MAPT and refeeding paradigm. Phosphopeptide abundance was normalized to group average protein abundance. P-values calculated by limma. Comparisons performed: WT vs. RI WF; WT vs. RII WF; within each line, WF vs. Stv 10’, WF vs. Stv 30’, and Stv 30’ vs. Refed 30’. Data are represented as mean ± SD. **(D)** Schematic illustrating known (PhosphoSitePlus) and predicted (NetPhos) PKA sites within Tau441. **(E)** Pearson’s correlational matrix between PKA subunit isoform abundances and pathology features, from human brain proteomics dataset in Johnson, et al. (2022). MAPT: total Tau abundance, measured by MS. MTBR and NonMTBR: abundance of peptides uniquely from Tau microtubule binding regions or not, respectively. NFT: standard Tau pathology metric. MMSE: mini-mental state exam; lower scores correspond to increased impairment. Diagnosis: 0, no AD; 1, AD. Braak: higher scores correspond to increased pathology. **(F)** MS quantification of MAPT PKA phosphosite pS673, corresponding to pS356 in Tau441, from human brain phosphoproteomics dataset in Ping, et al. (2020). CTL: control; ASYM: asymptomatic; AD: diagnosed AD. **(G-H)** Motif enrichment analysis of differentially abundant phosphopeptides (by p-value) in ASYM vs. CTL **(G)** and AD vs. CTL (H) binary comparisons. Only PKA motifs are shown. Cut-off for significantly enriched motifs was set to p < 0.00001 and minimum number of sequences per motif to 20.

Tau has 78 known (demonstrated by either low- or high-throughput, i.e., MS, methods) and 5 additional predicted (via NetPhos) phosphosites [3, 29]. PKA-C is an empirically demonstrated kinase for 14 of these known sites and is predicted to phosphorylate one additional site (prediction score 0.633, **Fig. 4D**) [29]. To begin to understand if dysregulation in PKA-R subunits is relevant to human disease, we took advantage of two rich MS datasets available via the Alzheimer’s disease (AD) Knowledge portal: a proteomics dataset consisting of over 1,000 brain tissues and a phosphoproteomics dataset of 27 brain tissues enriched for phosphopeptides [36, 37]. We hypothesized that PKA-related proteomic and phosphoproteomic changes in a human AD brain, compared with control patients would mirror the RI KO model. We performed pairwise correlational analyses between PKA subunit abundances, Tau abundance (whole MAPT protein, microtubule-binding domain (MTBR) peptides, and nonMTBR peptides), and patient disease data including diagnosis (0, no disease; 1, disease), Braak staging (0-6), mini mental status exam (MMSE) scores (0-30), and NFT pathology (**Fig. 4E**). RIα and RIβ abundance were positively correlated with one another and negatively correlated with RIIα, RIIβ, Cα, and Cβ abundance. RIα was also positively correlated with diagnosis stage and negatively correlated with MMSE score, whereas RIβ was negatively correlated with diagnosis, Braak stage, NFT pathology, and nonMTBR abundance. Most of these results align with the model that decreased abundance of Type I subunits, especially RIβ, corresponds to worsened disease states. On the other hand, RIIα, RIIβ, Cα, and Cβ abundance were all positively correlated with one another, as well as positively correlated to MTBR and total MAPT abundance. Cβ was additionally positively correlated to NFT pathology. Indeed, the RI KO cells closely mirror the proteomic changes with advancing disease: decreased Type I R subunits, increased Type II R and C subunits, and increased toxic Tau species. These findings are consistent with previous results in human brain tissue that link RII, Cβ, and NFTs [38, 39].

Phosphorylation of Tau at S356 has been reported in both prefibrillar and fibrillar tangles and is thought to stabilize Tau, making it resistant to degradation [40–43]. In the phosphoproteomics data from Ping, et al. (2020), we also found that Tau pS356 was significantly increased in AD brains, albeit only in symptomatic AD (**Fig. 4F**). Lastly, we searched this dataset for signatures of dysregulated PKA signaling via motif enrichment analysis using an analogous approach to **Fig. 3F** [32]. Two PKA motifs (RxRxxSp and RxxSp) were significantly altered in asymptomatic patient brains compared to non-AD controls, and ten motifs were significantly altered in AD patient brains, with clear overrepresentation of PKA motifs upregulated in AD (all 10 motifs, as opposed to 5 downregulated) (**Fig. 4G-H**). These findings illustrate that PKA catalytic activity is dysregulated in ADRD and may be a feature exclusive to later stages of disease, when R subunit regulation of PKA-C is lost.

### Tau shows preferential binding to PKA-RII isoforms over RI isoforms

Commensurate changes in PKA-C activity may alone explain the observation that loss of PKA-RI increases Tau phosphorylation and loss of RII decreases Tau phosphorylation. However, given that the R subunits not only regulate PKA-C activity but also define its subcellular localization through R:AKAP binding, we investigated if direct PKA:Tau interaction contributes to differential phosphorylation of Tau. A prior APEX2-based Tau proximity biotinylation study compared the proteins that bind the N-terminus of Tau to those that bind the C-terminus and found that RIIα and RIIβ interact with both termini of Tau, whereas RIα and RIβ only interact with the C-terminus (**Fig. S5A**) [44]. Another, BioID2-based proximity biotinylation study, only detected RIIβ-Tau interaction [45]. We saw a similar enrichment for RIIβ:Tau interaction (as opposed to tubulin) in our previous APEX-based Tau interactomics study as well (**Fig. S5B**). We therefore hypothesized that Tau binds more strongly to RII subunits than to RI subunits, thereby increasing RII holoenzyme localization to Tau.

We performed a peptide overlay analysis to identify the interacting regions within both the R subunits and Tau. First, the entire sequences of human RIα, RIβ, RIIα, and RIIβ were arrayed as 18-mer peptides staggered by three residues, then overlaid with either full-length (FL) Tau441 or Tau RD (K18) (**Fig. 5A-F, S6**). A human Tau-specific antibody (HT7) was used to detect bound Tau. Within each R-type, the binding patterns with Tau were similar; however, RIβ showed slightly higher signal compared to RIα, and Tau was able to bind the CNB-B domain of RIIβ but not RIIα (**Fig. 5A-F, SA-B**). Differences between the R-types were more striking: while Tau bound similar regions across subunits, Tau bound more strongly to RII in the linker region and CNB-A domain of RII and to RI, especially RIβ, in its CNB-B domain (**Fig. 5A-F, S6A-B**). There was minimal binding of Tau to either D/D domain, which suggests that Tau may not be a canonical AKAP, unlike its protein family member MAP2 [46, 47]. The binding patterns were largely reproduced in the peptide arrays overlaid with RD Tau, demonstrating that the RD alone is sufficient for R-Tau binding (**Fig. S6C-D**).

**Figure 5.**
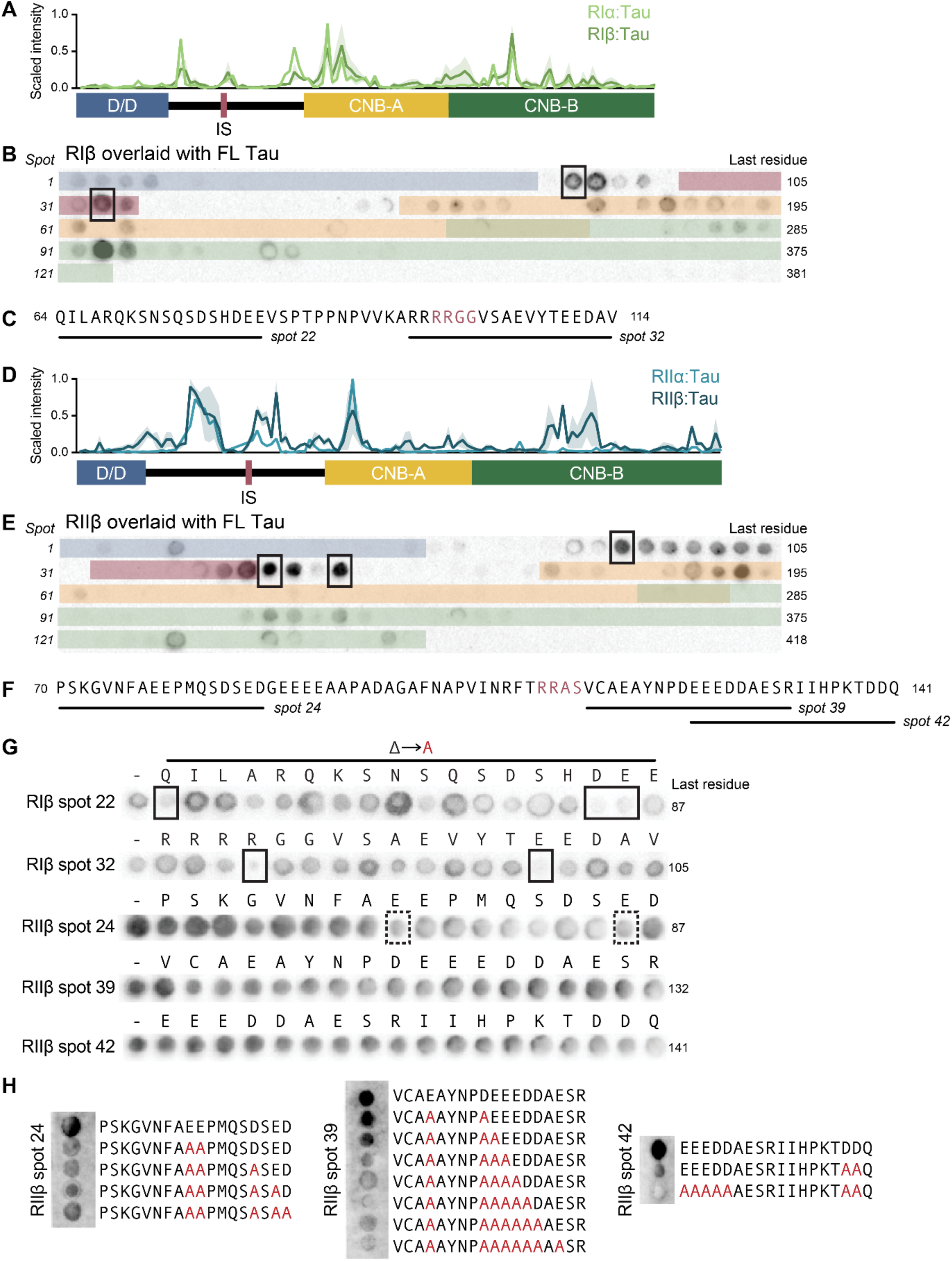
PKA demonstrates R-isoform-dependent preferential Tau binding. **(A-C)** Line graph of min-max scaled relative intensity (A) of RI peptide-binding arrays overlaid with FL Tau, corresponding to RIβ (B) and RIα (**Fig. S6**). RIβ peptide array was repeated twice, and the mean ± SEM is depicted by the shaded area. Domains in 5A and 5B correspond by color. Spots of interest in 5B are mapped onto the sequence of RIβ (C). **(D-F)** Line graph of min-max scaled relative intensity (D) of RII peptide-binding arrays overlaid with FL Tau, corresponding to RIIβ (E) and RIIα (**Fig. S6**). RIIβ peptide array was repeated twice, and the mean ± SEM is depicted by the shaded area. Domains in 5D and 5E correspond by color. Spots of interest in 5E are mapped onto the sequence of RIIβ (F). **(G)** Single-residue Ala scans of spots of interest from 5C and 5F, overlaid with FL Tau. (-) represents the original peptide, and the letter above each spot represents the residue within that peptide that was mutated to Ala. Spots in solid boxes represent those with >50% decrease of Tau:peptide binding, and spots with dotted boxes represent those with a possible decrease in Tau:peptide binding. **(H)** Multi-residue Ala scan for peptide spots of interest for RIIβ. Glu and Asp residues were mutated to Ala as shown in red.

We focused the remainder of our investigation on the RIβ and RIIβ subunits, as both are highly expressed in the brain and enriched in neurons as opposed to most other cell types (**Fig. S1B**). We further investigated Tau’s preferential binding to the R-linker region which is structurally divergent across R-types. Both RI and RII have flexible, disordered linker regions, but the RII-linker is richer in negative charges, more solvent exposed, and especially in the case of RIIβ, longer. To understand if a specific residue property drives Tau:R-linker binding, we performed Ala scans of the most prominent 18-mer spots within the RIβ and RIIβ peptide arrays, specifically RIβ spots 22 and 32 and RIIβ spots 24, 39, and 42 (boxed spots in **Fig. 5B**, **5E**). Mutating negatively-charged residues Asp and Glu to Ala diminished Tau binding to RIβ, but the Tau:RIIβ interaction was resistant to single residue mutations (**Fig. 5G**). Only when we performed poly-Ala scans mutating multiple Asp and Glu residues did we see diminished Tau:RIIβ binding, and only in spot 42 was the binding fully abolished, after all negatively-charged residues were mutated to Ala (**Fig. 5H**).

In summary, we found that Tau preferentially binds to PKA RII subunits over RI, especially within the flexible linkers, and this binding is dependent on the negatively-charged residues Asp and Glu. Our overall working model is that when RI is knocked out, the remaining RII_2_:C_2_ has a stronger interaction with Tau compared to WT holoenzyme, driven by electrostatic interaction with the RII-linker, increasing PKA-C localization to Tau and thereby Tau phosphorylation by PKA (**Fig. 6A**). Conversely, when RII is knocked out, the remaining RI has a weaker interaction with Tau compared to WT holoenzyme, leading to a decrease overall in Tau phosphorylation by PKA (**Fig. 6B**).

**Figure 6.**
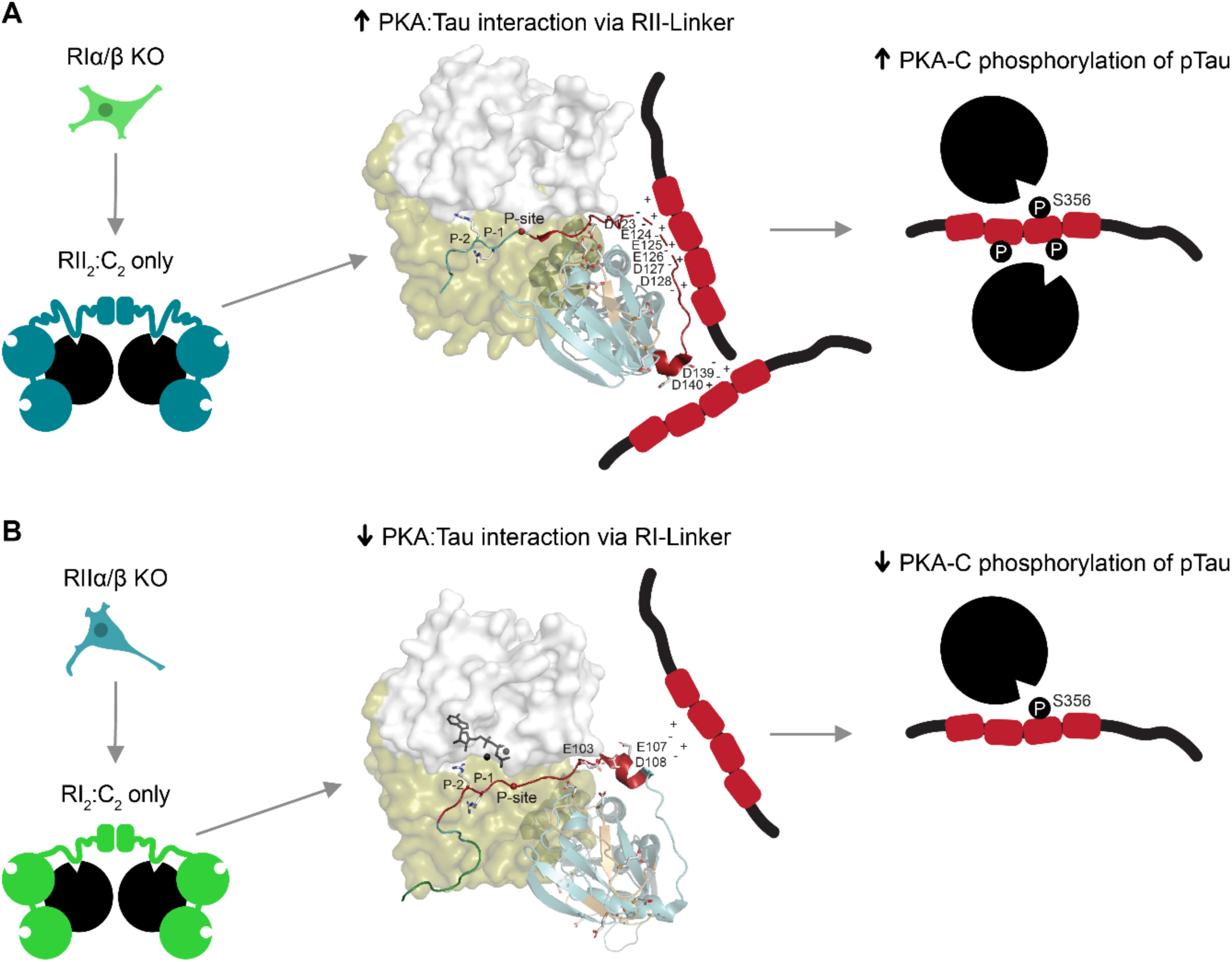
Working model of PKA-R-dependent regulation of Tau phosphorylation. **(A)** Model of increased PKA-driven phosphorylation of Tau in RIα/β KO cells. Schematic includes structure of mouse RIIβ:bovine C (PDB 3TNP), highlighting residues of interest within the linker region from peptide array (spots 39 and 42 from Fig. 5). **(B)** Model of decreased PKA-driven phosphorylation of Tau in RIIα/β KO cells. Schematic includes structure of human RIβ:bovine C with ATP and Mg^2+^ (PDB 4DIN), highlighting residues of interest within the linker region from peptide array (spot 32 from Fig. 5).

## DISCUSSION

In this study, we uncovered that loss of PKA Type I (RIα/β) vs. Type II (RIIα/β) R subunits leads to distinct, and often opposite, changes in the cellular proteome, phosphoproteome, and function. Consistent with earlier reports that RIα is the tissue-specific extinguisher of cAMP-mediated gene expression [48, 49], we found that loss of RI subunits led to an increase in Cα subunit abundance and kinase activity. Previous studies have demonstrated that, of the R subunits, only RIα was lethal when knocked out, and that additional KO of Cα and Cβ abolished the lethality of RIα loss, suggesting RIα alone is sufficient and necessary to regulate PKA activity [17]. In contrast, we found that loss of RII led to decreased Cβ abundance and kinase activity, consistent with previous literature showing that RIIβ is a key regulator specifically of Cβ. Using cutting-edge, MS-enabled omics, the present work builds upon previous literature in describing the functional nonredundancy of the R subunits in modulating PKA-C abundance and activity. Our results do not provide a clear answer on why the loss of RI vs. RII alters C subunit abundance. We provide evidence that this change is independent of Cα pS339, as we saw increasing Cα levels despite decreased pS339 levels which, based on previous knowledge, should have led to insoluble and nonfunctional Cα, the opposite of what we saw. We hypothesize that the RII_2_:C_2_ holoenzyme structure is more stable than the RI_2_:C_2_ complex, and future studies should explore the structural stability of various holoenzymes.

Proteomic and phosphoproteomic differences between RI and RII KO cells extend beyond basal conditions: a particularly striking aspect of our study is that loss of either RI or RII blunts metabolic flexibility in response to glucose starvation, which is a key function of intact PKA. Many metabolic PKA substrates, including PFK-2 and ACLY, showed baseline differences in phosphorylation, which typically followed the overall trend of PKA-C substrates, with RI loss increasing substrate phosphorylation and RII loss decreasing substrate phosphorylation. RI KO cells seemed particularly insensitive to glucose perturbations, with PKA substrate phosphorylation starting and staying high compared to WT, whereas the response in the RII KO cells was largely preserved. Notably, while our *in vitro* study does not model the physiological consequences of glucose starvation and the role of insulin, glucagon, and other hormones in the regulation of metabolism, our experimental conditions provide a reductionist perspective to elucidate acute, cell-autonomous signaling consequences of each R subunit family. Overall, our findings support a model in which different PKA-R subunits have distinct roles in the regulation of cellular metabolic flexibility and responses to nutrient stress and stimuli.

PKA, while one of the earliest reported Tau kinases, has been understudied and underappreciated for its role in Tau hyperphosphorylation and ADRD. It was serendipitous that the top differentially regulated phosphosite between RI and RII KOs was Tau pS356, underscoring the discovery potential of untargeted, MS-based proteomics and phosphoproteomics. In line with the changes we saw in many other PKA-C substrates, RI KOs had increased Tau pS356 compared to WT, whereas RII KOs had decreased Tau pS356. Taking advantage of two rich public datasets from brain tissue via the AD Knowledge Portal, we highlight the disease relevance of Tau pS356 and show that PKA subunit and activity changes in RI KOs closely resemble late-stage AD, positioning PKA as a therapeutic target for ADRD [36, 37]. PKA’s modulation of ADRD progression may extend further than Tau hyperphosphorylation, and we speculate that PKA has a significant role in integrating brain metabolism and synaptic function as well. Indeed, alongside synapse loss and Tau hyperphosphorylation and aggregation, metabolic dysfunction, including glucose hypometabolism, is a nascent hallmark of ADRD [50]. Additionally, metabolic syndromes associated with glucose metabolism, such as Type 2 diabetes, elevate AD risk [51]. Even if not causal, PKA is likely responding to glucose dysfunction in ADRD, and we showed in this study that dysregulation of PKA-R subunits further impairs the cell’s ability to modulate Tau phosphorylation in response to acute glycolytic stress. Metabolic interventions that alter glucose intake and metabolism, including ketogenic diets, caloric restriction, and intermittent fasting, are currently being evaluated in ADRD, and may already exploit PKA as a therapeutic target [52, 53]. For example, one study reported that the ketogenic diet, when administered later in life, benefits age-associated memory decline via PKA signaling in the presynapse [54].

Using peptide-binding arrays, we determined the interacting regions within both Tau and the R subunits, with the RII-linker showing preferential binding affinity for Tau over RI. Our results support a novel mechanism by which R subunits regulate PKA-driven phosphorylation of Tau: loss of RI increases Tau phosphorylation via increased C subunit abundance and activity and strengthened interaction of Tau with the holoenzyme’s remaining R subunit, RII. Understanding specific mechanisms by which PKA exacerbates Tau pathology allows for the development of therapeutics that modulate a specific aberrant function of a kinase in disease, for example RII:Tau binding, as opposed to the broad effects of drugs or metabolic interventions that nonspecifically target kinase activity. Lastly, future studies should seek to understand how Tau hyperphosphorylation impacts PKA-Tau binding, as each successive phosphorylation would decrease the overall negative charge of the MTBD, thereby altering its affinity for PKA-R’s negatively-charged linker.

A major strength of this study is the integration of proteomics, phosphoproteomics, and functional readouts. Phosphoproteomics remains a crucial tool for uncovering novel kinase biology, as we have with PKA-R and Tau, identifying novel phosphorylation sites, and validating kinase biology previously demonstrated through low-throughput methods. Limitations of this work, as with any MS study, include proteome coverage and stochasticity of detecting lower-abundance peptides and modifications. For example, key substrates regulated by PKA during starvation, including glycogen synthase, were not detected, therefore requiring pathway inferences without the full coverage of that pathway. Similarly, as we took an untargeted phospho-enrichment approach, we did not achieve full coverage of the PKA substratome, which should be expanded upon in further studies, such as using the RRxSp/Tp pan-PKA antibody as bait. Additionally, while peptide arrays have the advantage of being able to pinpoint specific interacting residues, they work best for identifying electrostatic interactions. Subsequently, we may be missing additional regions of PKA-R-Tau binding, including in PKA-R’s D/D domain, interactions with which may depend on its tertiary structure. However, we do not expect Tau to bind to the D/D domain, despite this being how R binds to the AKAP-like family member MAP2, as the D/D domain interacts with MAP2 on its N-terminus, whereas MAPs share a similar interaction in their C-termini. We also cannot detect protein interactions dependent on R_2_ or C_2_:R_2_ quaternary structures using this type of assay. Future work should seek to validate our current findings and identify novel sites of PKA-Tau interaction using the apo and holoenzyme proteins. Lastly, future work should investigate the convergent and divergent roles of all four isoforms, as while our work addresses overall differences between Type I and II R subunits, we also expect differences between each of their α and β isoforms.

## SUPPLEMENTAL FIGURES

**Figure S1.**
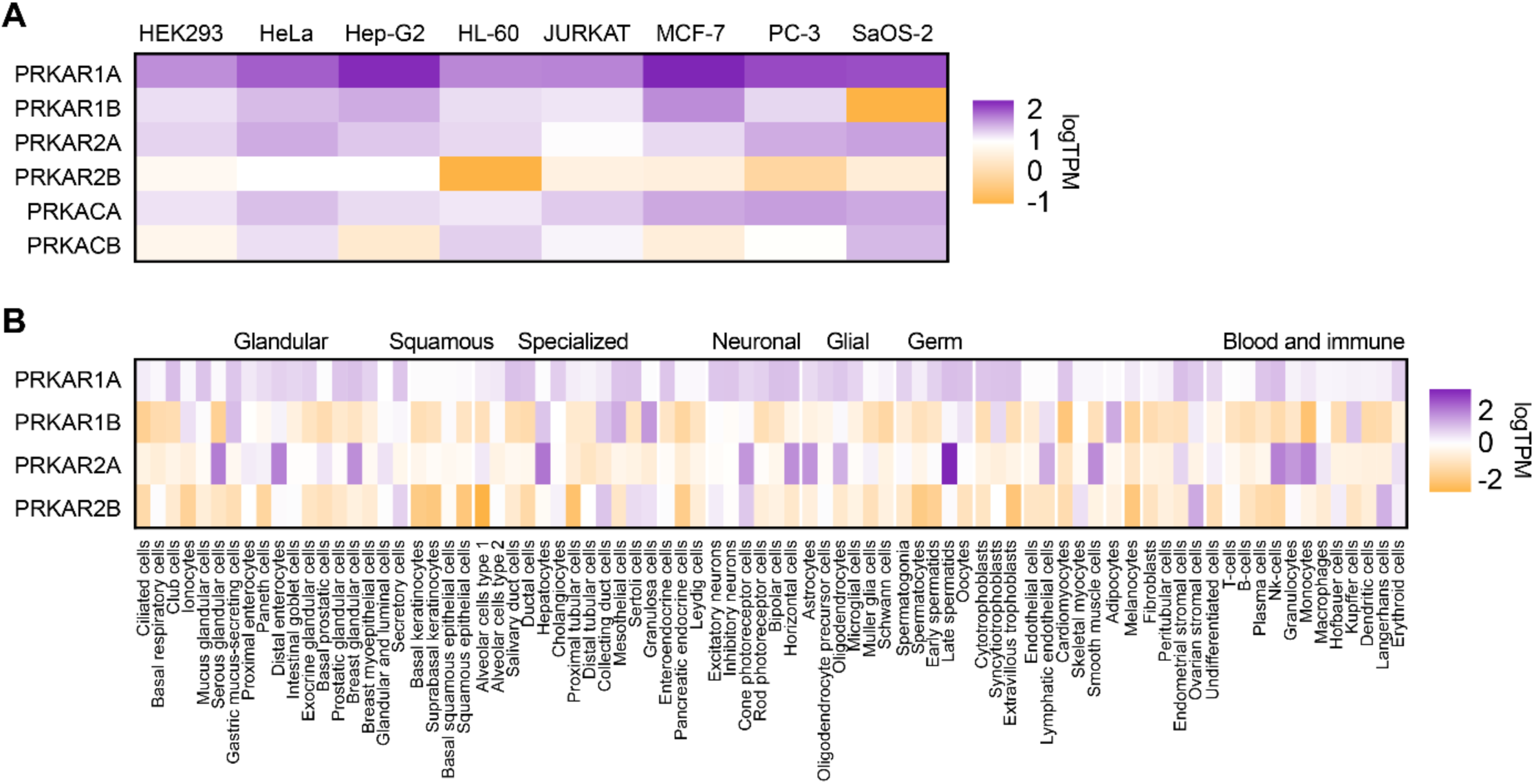
PKA subunit isoforms transcript counts across cell lines and types. Transcript counts for PKA subunit isoforms across cell lines **(A)** and cell types **(B)**, from Jin, et al. (2023) and Karlsson, et al. (2021), respectively.

**Figure S2.**
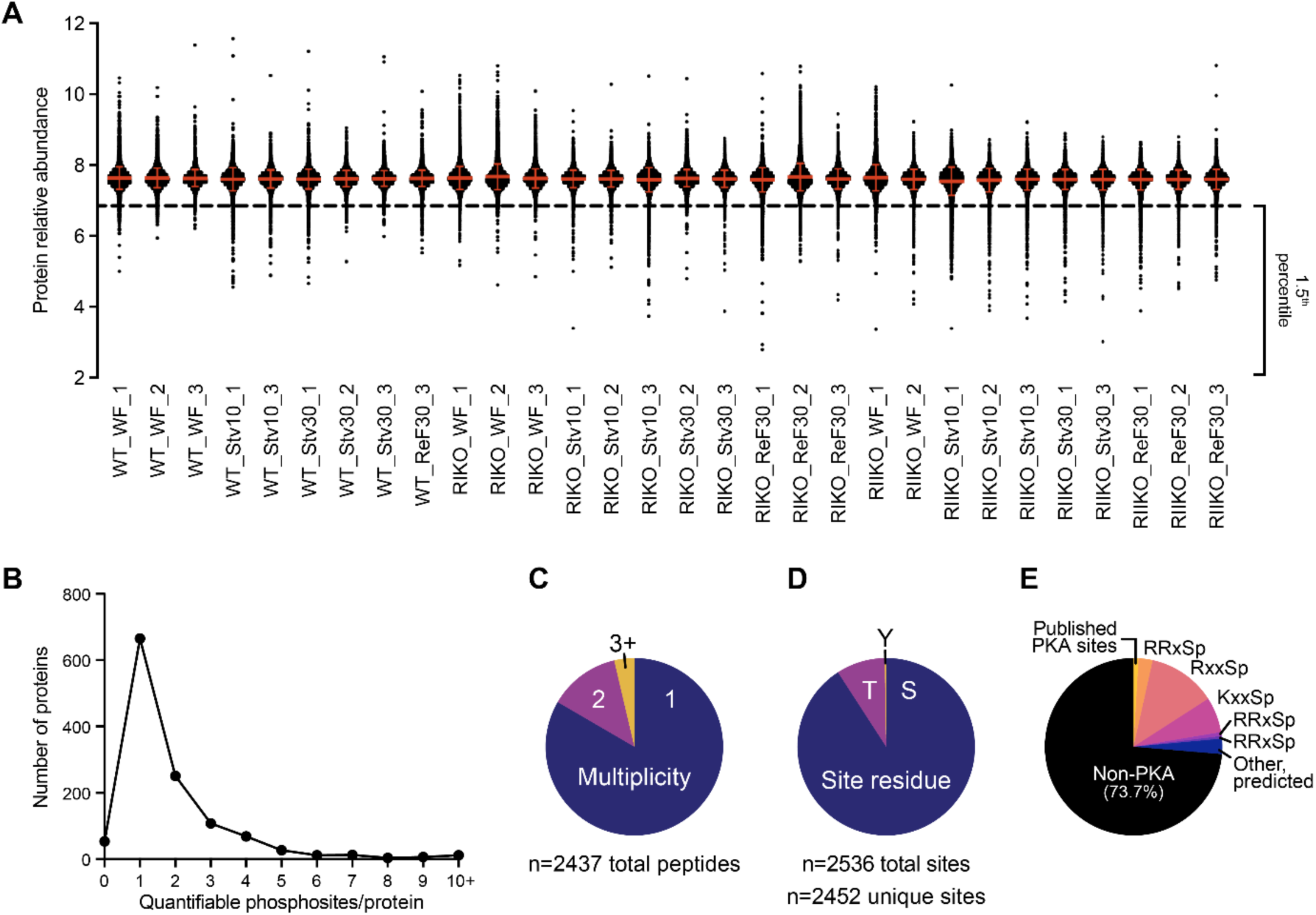
Supporting results from proteomics and phosphoproteomics study. **(A)** Relative protein abundances for all detected proteins in the present study, showing no total proteome normalization differences across plexes, groups, and replicates. y=6.88 line represents the positive detection threshold for this study, set at the 1.5^th^ percentile of all proteins detected. **(B)** Histogram illustrating the distribution of proteins represented in our phosphoproteomics study. Some phosphopeptides were identified (MS2) but not quantified (MS3). **(C-E)** Pie charts depicting proportions of phosphopeptides by feature detected in our study: multiplicity (C), residue type (D), and PKA motifs (E)

**Figure S3.**
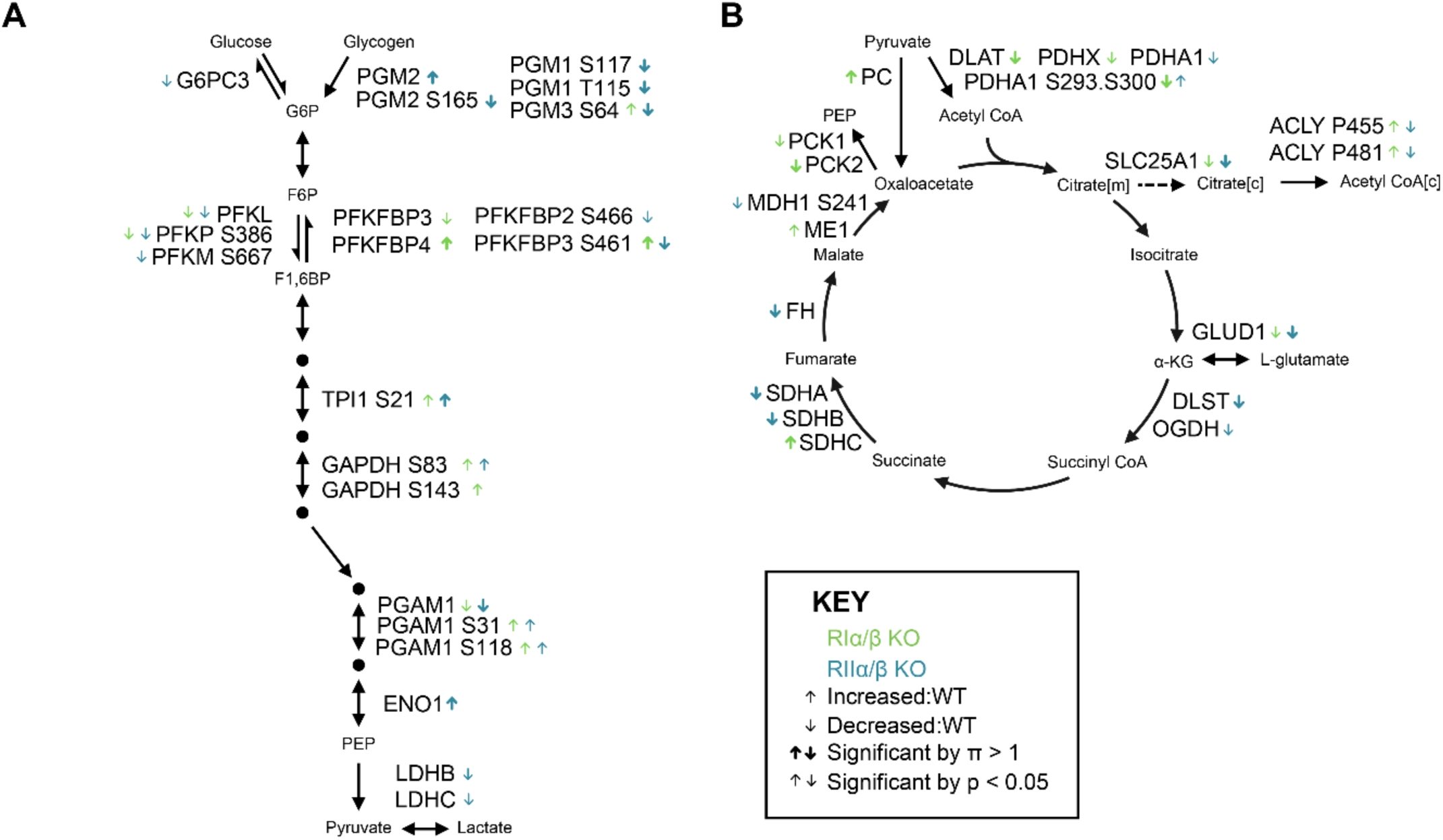
PKA RI and RII KO cells show distinct dysregulation of proteins and phosphosites regulating glucose metabolism. Pathway schematics of glycolysis/gluconeogenesis **(A)** and the citric acid cycle **(B)**, with significant hits from either RI KO vs. WT or RII KO vs. WT annotated. Directionality of the change is indicated by arrows. P-values were calculated by limma. Significant hits were defined as p < 0.05 only (thin arrows) or both p-value (<0.05) and π-score (>1) (thick arrows). Proteins or phosphosites not annotated were either not significant for either comparison or not detected in the present study.

**Figure S4.**
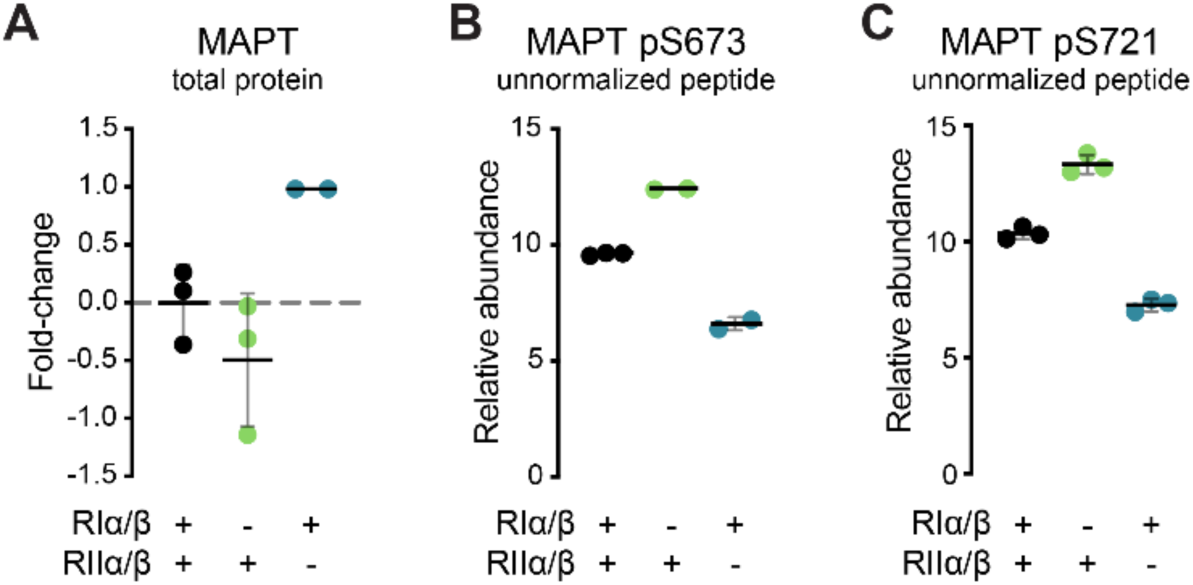
Corresponding to Fig. 5. Alterations in Tau phosphorylation are not due to total MAPT protein-level changes. **(A)** MS quantification of MAPT total protein by KO cell line, relative to WT. **(B-C)** MS quantification of MAPT phosphosite pS673 (B) and MAPT phosphosite pS721 (C), by KO cell line, relative to WT, corresponding to Fig. 4A**-B**. Raw peptide abundances not normalized to total protein level are represented. Data are represented as mean ± SD.

**Figure S5.**
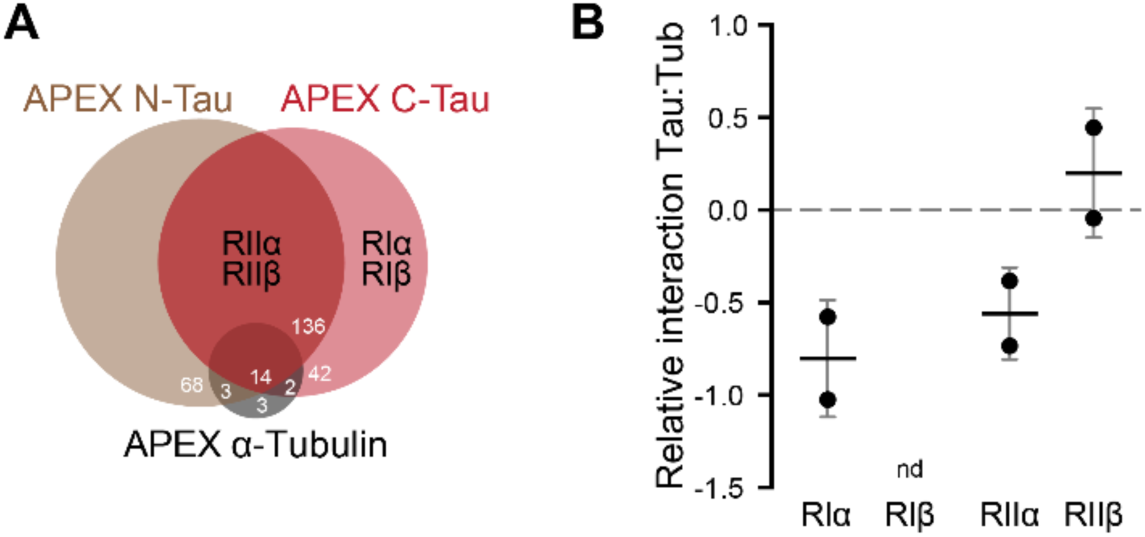
Corresponding to Fig. 5. APEX-Tau iPSC-neuron studies suggest strong Tau:RII interaction. **(A)** Venn diagram illustrating RI vs. RII interactions with Tau and tubulin, from study Tracy, Madero-Pérez, and Swaney, et al. *Cell* (2022). Labeling was done in iPSC-neurons, quantification was done by label-free MS. **(B)** R:Tau interaction relative to R:tubulin (Tub) from study Rossitto, et al. *BioRxiv* (2026). Labeling was done in iPSC-neurons, quantification was done by multiplexed MS.

**Figure S6.**
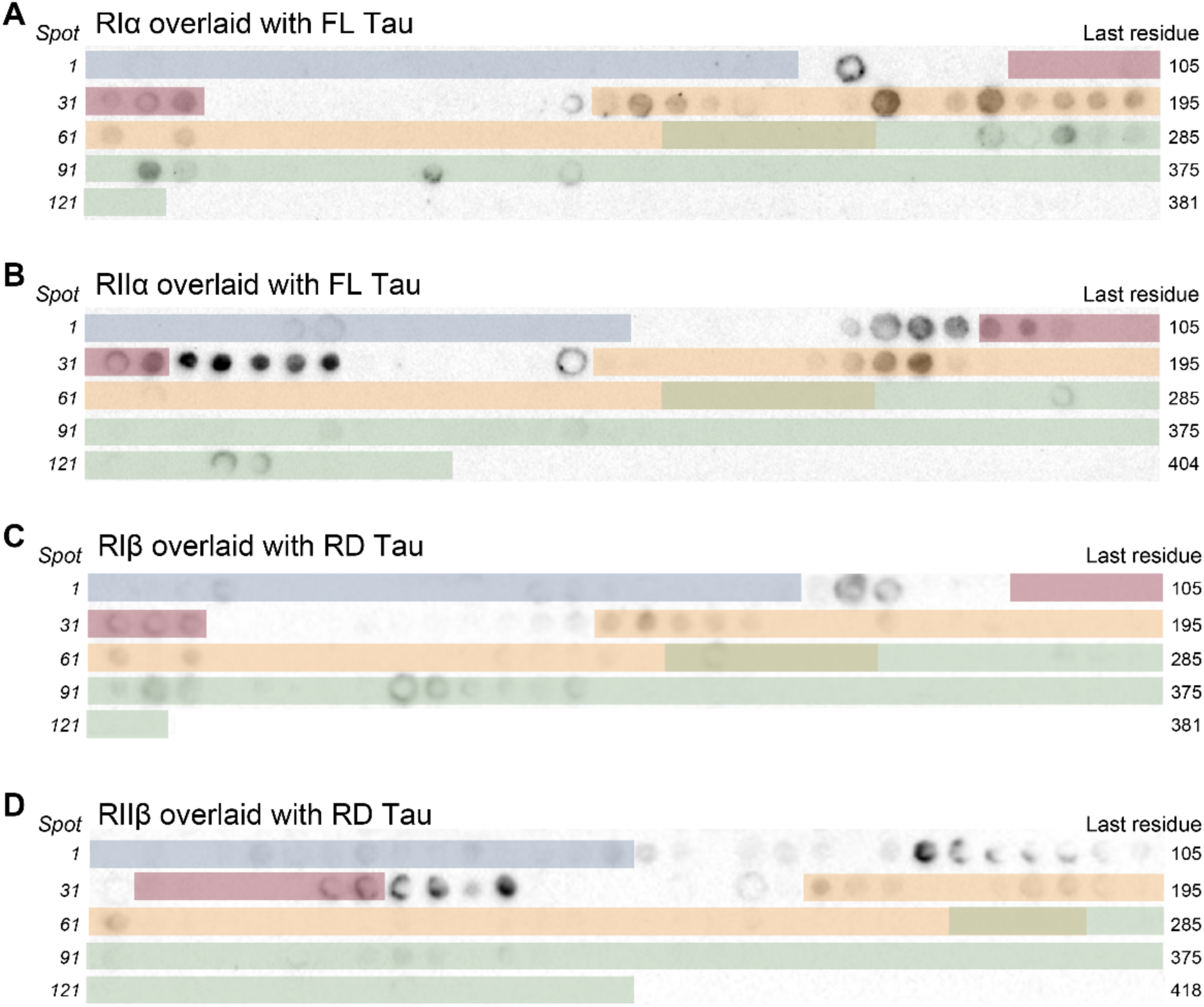
Corresponding to Fig. 6. PKA-R:Tau peptide arrays show strong RII:RD Tau interaction. **(A)** RIα peptide-binding array overlaid with FL Tau, quantified in Fig. 5A. **(B)** RIIα peptide-binding array overlaid with FL Tau, quantified in Fig. 5D. **(C)** RIβ peptide-binding array overlaid with RD Tau. **(D)** RIIβ peptide-binding array overlaid with RD Tau.

## MATERIALS & METHODS

### *In vitro* experiments

#### Generation of PKA-R double-KO cell lines

To interrogate the functional differences in cells with selective knockout of either PKA RI or RII subunits, CRISPR-Cas9 technology was used to edit human embryonic kidney (HEK293T) cells. HEK cells were chosen specifically because they are one of few cell lines that express all four R subunits, allowing us to compare the knockouts directly in the same system (**Fig. S1A**). To generate four different PKA R-subunits knock-out cells, we designed 20 bp gRNAs that specifically target early exons of human PKA R-subunits, and two gRNA were selected for each R subunit to ensure KO of any potential splicing isoforms:

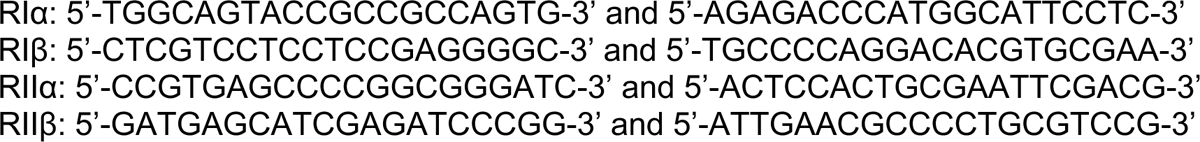

Each guide was cloned into the sgRNA scaffold of the PX458 backbone expressing SpCas9-2A-GFP (gift from Feng Zheng, Addgene #48138). Cells were transfected for 24 h, filtered through 35 μm strainers, and GFP+ cells were sorted using a BD FACSJazzTM Cell Sorter into single cells. Clones were validated by Sanger sequencing, WB, and MS (**Fig. 1C-E**).

#### HEK cell maintenance

WT, RI KO, and RII KO HEK293T cell lines were maintained in a cell culture incubator with 5% CO2 at 37°C. Unless otherwise specified for an experiment, cells were kept in complete DMEM media with 4.5 g/L *D*-Glucose, 110 mg/L sodium pyruvate, and 10% fetal bovine serum (FBS), which represents the basal or “well-fed” condition. For the MS experiment, 10M cells from each line were seeded into 15-cm dishes and allowed to grow to confluency, ∼20M cells/dish.

#### Starvation and refeeding paradigm

Cells were either left untreated (“well-fed”) or starved of glucose for 10’ or 30’. Glucose starvation was achieved by washing cells twice with PBS and adding warm DMEM with 10% FBS without glucose and sodium pyruvate (Invitrogen, #11966025). A subset of cells that were starved of glucose for 30 minutes were replenished (“refed”) with glucose (4.5 g/L) for 30’. This totaled to four conditions: well-fed, 10’ starved, 30’ starved, and 30’ starved and 30’ refed. Each condition was performed in triplicate, although due to the loss of one of the multiplexes during preparation, a few conditions only had two replicates for MS. At the end of each incubation, cells were washed twice with PBS and lysed with 2 mL of fresh lysis buffer (6M urea, 7% SDS, 50 mM TEAB, titrated to pH 8.1 with phosphoric acid) with added protease and phosphatase inhibitor tablets (Roche cOmplete protease inhibitor cocktail and PhosSTOP tablets at 1 mg/mL buffer). Lysates were sonicated for 3 x 5-sec rounds, and proteins were quantified by Pierce BCA Protein Assay Kit following manufacturer’s instructions.

#### Fluorescence resonance energy transfer (FRET) assay

PKA activity was measured using a FRET reporter assay based on AKAR4 [55]. The activity reporter contains a PKA substrate site and FHA1 domain flanked by two fluorescent proteins cpVenus (yellow) and Cireon (cyan) as a FRET pair (schematic in **Fig. 2E**). Channel and FRET intensities were recorded, and changes of the ratio between yellow and cyan correspond to PKA activity changes over the substrate. HEK cells were transfected with the reporter using Lipofectamine 3000 reagents (Thermo, #L3000008), following manufacturer’s protocol. Four hours prior to the FRET experiment, cells were switched into DMEM with no phenol red (for imaging) or glucose (to reduce ATP levels) (Invitrogen, #A1443001), with glutamine and 10% FBS. Shortly after FRET measurements began, either 4.5 g/L of glucose or galactose were added, and cells were recorded for an additional 10 min.

#### Cell stimulation

For drug treatments used in WB experiments, HEK293T cells were seeded in 6-well plates and serum-starved for 16 h. Cells were then stimulated with 1 μM forskolin (Sigma-Aldrich, #F6886) with 100 μM IBMX (Sigma-Aldrich, #I5879) for 1 h. For drug treatments used in CRE luciferase assays, HEK293T cells were serum-starved and stimulated with 1 μM forskolin, with or without 100 μM IBMX, for 16 h prior to the assay. Basal control cells received an equivalent volume of DMSO (vehicle).

#### CRE luciferase assay

HEK293T cells were seeded in 12-well plates and transfected with 500 ng of CRE-Firefly luciferase reporter plasmid (CRE-luc; Promega, #E8471) and 125 ng of a Renilla luciferase reporter vector (Promega) using TurboFect Transfection Reagent (Thermo Fisher Scientific, #R0531). The day after transfection, cells were serum-starved and treated with forskolin, with or without IBMX, as described above under Cell stimulation. Following treatment, luciferase activity was measured using the Dual-Glo Luciferase Assay System (Promega, #E2920) according to the manufacturer’s instructions. Luminescence was quantified using a Tecan Spark plate reader. Firefly luciferase values were normalized to Renilla luciferase values to control for transfection efficiency. The resulting values were then expressed relative to the basal WT condition.

#### Immunofluorescent microscopy

HEK cells were plated onto PDL-coated coverslips at equal densities and allowed to grow to 70-80% confluency, after which they were washed 1x with cold PBS and fixed with 4% PFA. Cells were then blocked in 10% BSA, stained for overnight then DAPI for 10 min, and mounted. A Nikon AXR confocal microscope system was used for imaging. All images were taken with the same confocal settings, including identical aperture and laser intensities, Galvano scanner settings, and number of scans averaging. Minor brightness and/or contrast adjustments were performed as necessary, consistently across all images in the same experiment. Antibodies and dilution used as follows: Cα (Taylor Lab in-house), Cβ (LSBio, #LS-C191947; 1:500), and AlexaFluor secondaries (ThermoFisher; 1:600).

### Biochemical experiments

#### Western blotting

Cells were lysed then sonicated in RIPA buffer containing 1:100 dilution PI, 1:100 dilution PIC2, 1:100 dilution PIC3, 1 mM PMSF, 0.5 mM NaF, and 1.6 nM TSA. Lysates were centrifuged at 16,000x*g* at 4°C for 15 min, and supernatants were collected. Protein concentrations were determined by the BCA assay (ThermoFisher, #23225). Equal amounts of protein were resolved on 4-12% Bis-Tris gels (ThermoFisher, NuPAGE system), transferred to either nitrocellulose or PVDF membranes depending on application, and probed with the specified antibodies. Total protein was visualized by Ponceau S stain. Bands in the immunoblots were detected using an enhanced chemiluminescence kit (ThermoFisher, #34094) and visualized using a ChemiDocMP Imaging System (Bio-Rad). Representative blots from the same membrane are shown. Antibodies and dilution used as follows: RIα (BD Transduction Laboratories, #610609; 1:500 with no milk) RIβ (R&D Systems, #AF4177; 1:1000), RIIα (SCBT, sc-137220; 1:200), RIIβ (Enzo, #BML-SA270; 1:1000), Cα (CST, #5842; 1:2000), Cβ (Abcepta, #AP7047a; 1:1000), p-RRXS*/T* PKA substrates (CST, #9624; 1:5000), pS133-CREB (CST, #9198; 1:2000), CREB (CST, #4820; 1:1000), HSP90 (CST, #4874; 1:5000), α-tubulin (CST, #3873; 1:7000), goat HRP-conjugated secondaries (Invitrogen, anti-Rb: #31460, anti-Ms: #31430; 1:2000), and donkey anti-Sheep HRP-conjugated secondary (R&D, #NBP1-73720; 1:2000).

#### Peptide arrays

Peptide spot arrays were generated by the INTAVIS AG peptide synthesizer (INTAVIS Bioanalytical Instruments AG, Koeln, Germany) using standard Fmoc (9-fluorenylmethoxycarbonyl) protection-based solid-phase peptide synthesis. 18-mer peptides staggered by three residues were directly conjugated onto amino-PEG modified cellulose membranes (INTAVIS AG, #ACS01). Membranes were activated via 5 min incubation with ethanol, blocked in 5% milk for two hours at room temp, then incubated with either Tau441 (rPeptide, #T-1001) or homemade RD Tau (i.e., K18-Tau) monomers in TBST (TBS, pH 8.0 with 0.1% Tween-20) overnight at 4°C. The following day, the array was washed 3x TBST then incubated with human Tau-specific antibody HT7 (Invitrogen, #MN1000; 1:1000) overnight at 4°C. The last day, the array was washed, incubated in goat anti-Ms HRP-conjugated secondary (Invitrogen, #31430), washed again, then developed using enhanced chemiluminescence kit (Thermo, #34094) and visualized using a ChemiDocMP Imaging System (Bio-Rad). Spot integrated density was measured in ImageJ, and spots within a single membrane were min-max scaled, with the min value representing a spot with no binding. Alanine scans were generated by replacing each residue with Ala one at a time and otherwise performed the same way as full scans.

### Quantitative multiplexed proteomics and phosphoproteomics

#### Peptide preparation

Lysates from the five starvation and refeeding conditions were analyzed by quantitative multiplexed proteomics, starting with approximately 20M cells and 5 mg of protein per lysate. Disulfide bonds were reduced in 5 mM dithiothreitol (DTT) at 47°C for 30 min, and free cysteines were alkylated in 15 mM iodoacetamide at room temperature, darkened environment for 20 min. The alkylation reaction was quenched for 15 minutes at room temperature through the addition of an equivalent volume of DTT as in the reduction reaction. Proteins were then trapped using S-Trap mini columns (ProtiFi, #C002-MINIX), digested with trypsin (Promega, #V5113), and eluted according to manufacturing protocols. Eluents were desalted on SepPak C18 columns using instructions provided by the manufacturer (Waters). Desalted samples were dried under a vacuum. Peptides were quantified using a Pierce Colorimetric Peptide Quantification Assay Kit following manufacturer’s instructions. 50 μg of each sample was separated for proteomic analysis (next step: labeling), and 4 mg was separated for phosphopeptide enrichment. Peptides were stored at -80°C.

#### Phosphopeptide enrichment

Phosphopeptides were enriched by TiO2 beads following previously established protocols. TiO2 beads were prepared by washing once with binding buffer (2 M lactic acid, 50% acetonitrile (ACN)), one with elution buffer (50 mM KH2PO4, pH 10), and twice again with binding buffer. Peptides were resuspended in binding buffer, combined with beads at a ratio of 1:4 peptides:beads, and vortexed at room temperature for one hour to allow phosphopeptides to bind beads. Beads were then washed three times with binding buffer and three times with wash buffer (50% ACN/0.1% trifluoroacetic acid (TFA)). Phosphopeptides were eluted from the beads by two 5-minute mixing incubations with elution buffer. Enriched peptides were desalted on SepPak C18 columns, dried under a vacuum, and stored at -80°C until labeling.

#### Labeling

Peptides for proteomic and phosphoproteomic analyses were labeled separately using Tandem Mass Tag (TMT) 10-plex reagents (lot number VA296083), each experiment with six 10-plexes. Peptides were resuspended in 50 μL of 30% ACN with 200 mM HEPES, pH 8.5. TMT labels were suspended in anhydrous ACN to a final concentration of 20 mg/mL, and 7 μL of the appropriate tag were added to each resuspended sample. For proteomic analysis, each 10-plex was “bridged” using a pooled control in channel 126, which consisted of equal peptide contributions from each sample to maximize the number of proteins identified and minimize missing values. Samples were randomly assigned to the remaining nine TMT labels for each experiment. For phosphoproteomic analysis, the samples were not bridged with a pooled control and were instead computationally bridged as later described (**Data processing and normalization**), to maximize the number of unique phosphosite identifications. The labeling schemes are included in the Supplemental Material. (Two treatment groups included in the MS experiment and therefore labeling scheme were not included in the final analyses, in one part due to a desire to simplify treatment groups for easier interpretability and the other due to proteomics plex #5 being compromised during fractionation and therefore not sequenced.) The labeling reaction proceeded at room temperature for one hour, after which excess label was quenched through the addition of 8 μL of 5% hydroxylamine for 15 min. 50 μL of 1% TFA was added to each sample, then samples were combined into the appropriate multiplexes. Plexes were desalted on SepPak C18 columns and dried under a vacuum.

#### Fractionation

Multiplexed samples were next subjected to fractionation using reverse phase high pH liquid chromatography to increase sequencing depth. An Ultimate 3000 high performance liquid chromatography system fitted with fraction collector, C18 column (4.6 x 250 mm), solvent degasser, and variable wavelength detector was used for fractionation. Samples were resuspended in 105 μL of 25 mM ammonium bicarbonate (ABC) and fractionated on a 22% to 35% ACN gradient with 10 mM ABC over an hour. The resulting 96 fractions were concatenated into 24 fractions by combining alternating wells within each column, and 12 alternating fractions were used for mass spectrometry analysis.

#### MS-based proteomic and phosphoproteomic analysis

Labeled peptides were resuspended in 5% ACN/5% formic acid and analyzed on an Orbitrap Fusion Tribrid mass spectrometer with an in-line Easy-nLC 1000 System. All data acquired were centroided. 1 μL of each multiplex was loaded onto a 30 cm in-house pulled and packed glass capillary column (I.D. 100 μm, O.D. 350 μm). The column was packed with 0.5 cm of 5 μm C4 resin followed by 0.5 cm of 3 μm C18 resin, with the remainder of the column packed with 1.8 μm of C18 resin. Electrospray ionization was assisted by the application of 2000 V of electricity through a T-junction connecting the column to the nLC, heating the column to 60°C. Following sample loading, peptides were eluted using a gradient ranging from 6% to 25% ACN with 0.125% formic acid over 165 minutes at a flow rate of 300 nL/min.

MS1 spectra were acquired in data-dependent mode with a scan rate of 600-1200 m/z for proteomics and 500-1500 m/z for phosphoproteomics and a mass resolution of 60,000. Automatic gain control (AGC) was set to 5e4. Maximum injection time was set to 35 ms for proteomics and 70 ms for phosphoproteomics. Lower ion intensity threshold was set to 5.0e4. Ions selected for MS2 analysis were isolated in the quadrupole at 0.5 Th. Ions were fragments using collision induced dissociation, with a normalized collision energy of 30% and were detected in the linear ion trap with a rapid scan rate. MS3 analysis was conducted using the synchronous precursor selection option to maximize TMT quantitation sensitivity. Up to 10 (proteomics) or 3 (phosphoproteomics) MS2 ions were simultaneously isolated and fragmented with high-energy collision-induced dissociation using a normalized energy of 55%. MS3 fragment ions were analyzed in the Orbitrap at a resolution of 60,000. The AGC was set to 5e4 using a maximum ion injection time of 100 ms for proteomics and 250 ms for phosphoproteomics. MS2 ions 40 m/z below and 15 m/z above the MS1 precursor ion were excluded from MS3 selection for proteomics and 20 m/z below and 15 m/z above were cutoffs for MS3 selection for phosphoproteomics.

#### Data processing and normalization

Raw mass spectrometry files were processed using Proteome Discoverer 2.5. MS2 data were queried against the Uniprot human database (downloaded May 2022) using the SEQUEST algorithm. A decoy search was also conducted with sequences in reverse order. Data were searched using a precursor mass tolerance of 50 ppm and fragment mass tolerance of 0.6. Static modifications were specified as follows: TMT 10-plex reagents on lysines and N-termini and carbamidomethylation of cysteines. Dynamic modifications were specified to include oxidation of methionines and, for the phosphoproteomic experiment, phosphorylation of serine, threonine, and tyrosine residues. The enzyme specificity was set to full trypsin digest with two missed cleavages permitted. Resulting peptide spectral matches were filtered at a 0.01 FDR by the Percolator module against the decoy database.

For the proteomics experiment, peptide spectral matches were exported and summed to the protein level, using only spectra with a signal-to-noise per label >10 and isolation interference of <25%. Unique peptides of all PKA isoforms were detected, allowing for accurate differentiation and quantification of the isoforms. Specifically, RIα was classified by 17 unique peptides, RIβ by 3 peptides, RIIα by 15 peptides, RIIβ by 10 peptides, Cα by 3 peptides, and Cβ by 2 peptides. Data were then batch corrected in a multistep process as previously described [56]. First, data are standardized to the average for each protein and then to the median of all averages for protein. Then, to account for slight differences in amounts of protein labeled, these values are then standardized to the median of the entire data set. Lastly, the data were normalized by log2 transformation. Final data represent normalized relative protein abundances. Proteins with relative abundances below the 1.5^th^ percentile of all detected proteins were considered below the positive detection threshold.

For the phosphoproteomics experiment, MS phosphopeptide quants were transformed into natural log scale and median-centered. Technical variation (a.k.a. “batch effects”) between the TMT 10-plexes was removed using linear modeling as implemented in the limma R library [57]. To assess batch correction quality, we hierarchically clustered the samples and monitored a decrease in the Adjusted Rand Index (ARI) representative of TMT batches. Lastly, phosphopeptide abundances were divided by the average protein abundance for their biological group to account for any abundance differences at the protein level while minimizing loss of data due to missing protein values in some plexes. Final data represent normalized relative phosphopeptide abundances unless otherwise specified.

#### Statistical analyses

Statistical tests were performed on normalized protein and phosphoprotein abundances, which for the proteomics data represent the log2 normalized abundances, and for the phosphoproteomics data represent the log2 normalized abundances divided by the normalized group average protein abundance. For the basal/well-fed (WF) experiment, two binary comparisons were performed: WT vs. RI KO and WT vs. RII KO. For the starvation and refeeding experiment, three binary comparisons were performed within each line: 10’ stv vs. WF, 30’ stv vs. WF, and 30’ refed vs. 30’ stv. P-values were calculated by limma using the lmFit and eBayes functions (R, *limma* 3.60.3), and the significance cut-off was p < 0.05 [57]. For untargeted analyses, such as those in Fig. 1, an additional, stringent π-score cut-off (-logP x |log2(fold-change)|) of > 1 was also used to correct for multiple hypotheses [58].

#### *K*-means clustering

*k*-means clustering was performed in R using the kmeans function (R, *stats* ver. 4.4.3) on the starvation and refeeding phosphoproteomics data. Only the WT line was used to train the algorithm, to illustrate how the KO cells respond to glycolytic stress compared to the “expected” WT response. The appropriate number of clusters was determined via the elbow method to be eight. Scaled abundances (within each peptide) were graphed with *pheatmap* (R, ver. 1.0.12).

#### Motif enrichment analyses

Logos for significantly increased or decreased phosphopeptides in binary comparisons were generated using WebLogo (web, ver. 3.7.4). Motif enrichment analysis was done using motifx (R, *rmotifx* ver. 1.0). Foreground sequences were sequences of total length 15, ± 7 amino acid residues flanking each phosphosite, that were either significantly increased or decreased in binary comparisons by p-value. Background sequences were extracted from the human proteome (Uniprot UP000000589) using helper packages parseDB and extract Background (R, *PTMphinder* ver. 0.1.0). The central residue was set to S or T, the minimum sequence cut-off was set to 20, and the p-value cut-off was set to 1x10^-5^. Enriched motifs are represented by their motif score, which is calculated by taking the sum of the negative log probabilities used to fix each position of the motif. Higher motif scores correspond to more specific and statistically significant motifs. Only PKA motifs were graphed.

#### Kinase prediction

Kinase prediction was performed in two ways, primarily via NetPhos (web, ver. 3.1), with positive predictions having scores >0.5. Additionally, CST’s PhosphoSitePlus (web) resource was used to identify other potential PKA-C sites detected in the study, with positive prediction thresholds set to either a top ten score or percentile rank or top 10^th^ percentile score for at least one PKA-C isoform. PhosphoSitePlus identified 71 additional predicted PKA sites beyond RxxSp/Tp and KxxSp/Tp, many of which were RxSp/Tp, KxSp/Tp, RSp/Tp, KSp/Tp, or downstream of histidine.

### Human brain MS analyses

MS data of human brains was obtained from the AD Knowledge Portal via the Synapse repository under an approved data use certificate. Proteomics dataset was generated by Johnson, et al. and is available under the Synapse repository ID syn25006611 [37]. Five MAPT isoforms were quantified in this study, and we performed our analyses using MAPT_P10636-6, as it was detected in all brains and had the highest normalized abundance on average. The phosphoproteomics dataset was generated by Ping, et al. and is available under the Synapse repository ID syn20820053 [36]. The original, published p-values from phosphoproteomics data were used for significance cut-offs, and phosphosites with p < 0.05 were used for motif enrichment analysis. Motif enrichment analysis was performed as described in ***Quantitative multiplexed proteomics and phosphoproteomics***, with the central residue set to S or T, the minimum sequence cut-off set to 20, and the p-value cut-off set to 1x10^-5^. Only PKA motifs were graphed.

### Data representation and statistical analysis

Each point on a graph represents an independent biological replicate. Graphs were made in GraphPad Prism (ver. 10) and R (ver. 4.4.3), and aesthetics were adjusted in Adobe Illustrator (2024). Mean ± SD are represented on all graphs unless otherwise specified in the figure legend. p < 0.05, p < 0.01, p < 0.001, p < 0.0001, and not significant represented as *, **, ***, ****, and ns, respectively, and represent pairwise comparisons. # symbols were used occasionally to show comparisons to WT within the same treatment, and here p < 0.05, p < 0.01, p < 0.001, and p < 0.0001 are represented as #, ##, ###, and ####, respectively. A diamond was used to show hits that are additionally significant by π-score, and ◆◆◆◆ represents p < 0.0001 and π > 1.

Statistical methods for the proteomics and phosphoproteomics studies are described in ***Quantitative multiplexed proteomics and phosphoproteomics***. Statistical tests and multiple comparisons were decided *a priori* and kept consistent throughout experimental designs. For two-variable experiments, full-model two-way ANOVA and Šídák’s multiple comparisons tests were performed. For correlational analyses, Pearson’s correlation was performed.

## Data availability

Raw spectral files can be found at MassIVE (MSV000096782) and ProteomeXchange (PXD059469). A workbook including spreadsheets of the raw and normalized protein and phosphoprotein quantifications, statistical tests, and lists of proteins/phosphoproteins used for each analysis and results is included in Supplemental Materials. Human brain data from published proteomics and phosphoproteomics data is available through the AD Knowledge Portal (Synapse repositories syn25006611 and syn20820053, respectively) with an approved data use certification. This paper does not report original code. This study did not generate new unique reagents. Requests for further information and resources should be directed to and will be fulfilled by the lead contacts, Susan Taylor and Xu Chen.

## ACKNOWLEDGEMENTS

This work was supported by National Institutes of Health grants R01AG074273 (X.C.), R01AG078185 (X.C.), T32GM007752 (L.M.R.), T32AG066596 (L.M.R.), F99AG088570 (L.M.R.), R35GM130389 (S.S.T.), and the Health Science Center for Proteomics at UCSD (D.J.G.). We thank C. Sanchez, Y. Tao, and Y. Liu, for technical assistance and members of the Chen, Taylor, and Gonzalez labs for discussions.

## AUTHORS CONTRIBUTIONS

Conceptualization, L.M.R., T.L., X.C., and S.S.T.; investigation, L.M.R., T.L., Y.M., P.K.S., V.B., C.C.G., J.B., S.M., J.W., A.L., and I.K.; writing—original draft, L.M.R., X.C., and S.S.T.; Writing—review & editing, all authors; visualization, L.M.R., T.L., V.B., P.K.S., and J.W.; funding acquisition, L.M.R., D.J.G., X.C., and S.S.T.; resources and supervision, I.K., J.S.G., D.J.G., X.C., and S.S.T.

## CONFLICTS OF INTEREST STATEMENT

The authors declare that they have no conflicts of interest.

## Notes

### Competing Interest Statement

The authors have declared no competing interest.

